# Multidimensional profiling of heterogeneous lateral habenula subpopulations reveals distinct responses during motivated behavior

**DOI:** 10.64898/2026.08.05.743065

**Authors:** Morgan B. Corniquel, Joseph M. Martinez, Lina M. Hinostroza, Julien Gonzalez-Palavicini, Sydney Spytek, Michael L. Wallace

**Affiliations:** Graduate Program for Neuroscience, Boston University, Boston, MA; Center for Systems Neuroscience, Boston University, Boston, MA; Center for Neurophotonics, Boston University, Boston, MA; Department of Anatomy & Neurobiology, Boston University Chobanian and Avedisian School of Medicine, Boston, MA 02118, USA

**Author notes:** These authors contributed equally (Morgan B. Corniquel, Joseph M. Martinez).

**Keywords:** habenula, reward prediction error, motivation

## Abstract

The lateral habenula (LHb) shapes reward and aversion learning via projections to midbrain monoaminergic centers. Recent studies have demonstrated significant genetic, anatomical, and electrophysiological diversity within the LHb. However, it remains unclear how genetic or intrinsic electrophysiological characteristics relate to *in vivo* neuronal activity patterns. Detailed descriptions of transgenic mouse lines labeling LHb are lacking, limiting genetic access to LHb neuronal subtypes. Here we describe spatial gene expression patterns, electrophysiological characteristics, and projection targets for specific subpopulations of neurons in the LHb targeted via existing transgenic mouse lines. Furthermore, we demonstrate that two genetically defined subpopulations differentially respond to value, prediction errors, and directional movement during flexible, reward-guided behavior. These findings indicate that specific, genetically targetable, neuronal subpopulations in LHb may control discrete aspects of motivated behavior through parallel circuits targeting serotonergic and dopaminergic midbrain centers.

## Introduction

To survive in dynamic environments, animals need to monitor action outcomes and adjust strategies accordingly. Neuronal circuits required for reward-based learning compute error signals denoting the difference between expected and observed outcomes, or reward prediction errors (RPE).^1–3^ RPE signals can be positive, indicating that the experienced outcomes were better than expected, or negative, meaning the outcome was either worse or more aversive than anticipated.^4^

A key brain area implicated in generating RPE signals is the lateral habenula (LHb). This epithalamic structure is phylogenetically conserved across vertebrates and has been studied in animals ranging from zebrafish through rodents to humans.^5–7^ Inputs to the LHb include, among others, the basal ganglia, basal forebrain, lateral hypothalamus, and prefrontal cortex.^8–10^ These regions carry diverse signals including ongoing action selection, behavioral state, and previous action outcomes which are integrated by the LHb to signal prediction errors to downstream structures, including the ventral tegmental area (VTA), rostromedial tegmental nucleus (RMTg), and raphe nuclei (RN).^8, 11–14^ Specifically, LHb activity conveys a negative reward prediction error (nRPE), increasing following the omission of an expected reward or the presentation of an unexpected aversive stimulus.^15^ The LHb, which consists of glutamatergic neurons, canonically inhibits downstream dopaminergic and serotonergic neurons by exciting intermediate GABAergic inhibitory neurons within the VTA, RN, or the RMTg.^12, 16–18^ However, glutamatergic LHb projections have also been shown to provide direct connections to downstream midbrain dopaminergic and serotonergic neurons, suggesting a complex circuitry and function for LHb outputs based on projection targets.^19–21^

LHb neurons display several distinct spontaneous action potential (AP) firing patterns which have been described as silent (no APs), tonic (regularly spaced APs), or bursting (bimodal inter-spike interval (ISI) distributions).^22, 23^ Hyperactivation of the LHb and an increase in APs firing in a burst pattern have been observed following chronic stress and correlate with behavioral changes, including anhedonia.^23, 24^ Behavioral phenotypes can be reversed through LHb ablation or through the reduction of burst pattern APs with pharmacologic manipulations such as ketamine.^16, 23, 25^ The activation and maintenance of AP bursts, as opposed to regular tonic APs, is dependent on activation of low-voltage sensitive T-type calcium channels, N-methyl-D-aspartate (NMDA) receptors, and hyperpolarization-activated cyclic nucleotide-gated (HCN) channels.^23, 26, 27^ While roughly 10% of neurons appear to spontaneously fire AP bursts, a greater percentage can be driven to produce AP bursts after the release of a brief hyperpolarizing current, termed rebound bursting.^23, 24, 28^ While the propensity of a neuron to produce APs in a burst pattern is likely due to differential gene expression of the underlying channels, there is a lack of available tools to target bursting LHb neurons *in vivo*, hindering specific manipulation and observation.

LHb control over downstream structures and behavior is likely mediated by neuronal subpopulations that differ in their spatial location, input structures, downstream projection targets, and gene expression.^29^ Subpopulations have been described using cellular morphology, axonal tracing, and single-cell transcriptomic sequencing (sc-seq).^30–34^ Studies have broadly segmented the LHb into subnuclei and have identified differential gene expression patterns including genes required for burst-firing AP patterns.^32^ It is unclear, however, whether these genetically defined neuronal subtypes distinctly impact behavior or are differentially incorporated into expanded neural circuits.

In this study, we connect LHb spatial gene expression patterns, intrinsic electrophysiological characteristics, and projection targets for LHb neurons targeted using transgenic mouse lines. Further, we show differential LHb subpopulation activity in flexible, reward-guided behavior in a freely moving, probabilistic switching task exploring subpopulation activity in relation to reward, reward history, and direction of movement during the task. Detailed 3D mapping of axonal projections and recordings of neuronal activity across neuronal subclasses of the LHb provides insight into how these neurons differentially signal expectation, choice, and outcome as well as how they may differentially modulate monoaminergic signaling in downstream regions. Together we provide detailed analysis of genetically targetable subpopulations within the LHb which show temporally and functionally distinct activity patterns during motivated behavior.

## Results

### Transgenic mouse lines target genetically defined LHb neuronal subpopulations

Sc-seq studies have described genetically distinguishable neuronal subpopulations in the LHb.^32–34^ However, the spatial distribution of these subpopulations, particularly along the rostral/caudal axis of the LHb, is unknown. We chose four genes (*Sst, Peg10, Rbfox1*, and *Vgf*) which we and others have shown to label genetically distinct subclasses using sc-seq.^32–34^ We performed multiplexed fluorescent in-situ hybridization (FISH) to quantify the expression and spatial distribution of these four transcripts throughout the LHb (Fig. S1A-C). Consistent with previous results, we found that expression patterns of these transcripts were largely non-overlapping, occupying different regions of the LHb (Fig. S1D-K).^32^ *Sst* was expressed in a discrete region of the dorsal/posterior LHb and likely overlapped with the HbX subregion (Fig. S1D-E).^35, 36^ *Peg10* expression largely occupied the center of the medial/lateral extent of the LHb, within the posterior half of the anterior/posterior axis (Fig. S1F-G). Unlike *Sst* and *Peg10*, *Rbfox1* and *Vgf* showed considerable overlap in individual neurons, predominantly in the anterior pole of the LHb (Fig. S1C, S1H-K). In the remaining anterior/posterior extent, *Vgf* showed greatest expression levels in the most medial portions of the LHb, while *Rbfox1* was expressed in both the most medial and lateral fringes of the LHb, flanking *Peg10* expression (Fig. S1H-K). Therefore, detailed spatial analysis across the anterior/posterior extent of the LHb revealed that *Sst, Peg10, Rbfox1*, and *Vgf* are largely expressed in distinct LHb neurons, with the notable exception of high levels *Vgf* and *Rbfox1* in individual neurons at the anterior pole of the LHb.

Following queries of published sc-seq and *in situ* gene expression data,^32, 37^ we hypothesized that three Cre-recombinase expressing transgenic mouse-lines (*Sst*-Cre,^38^ *Kcnc1*-Cre,^39^ and *Prokr2*-Cre), would show subpopulation specific expression patterns in LHb. To genetically profile *Cre* expressing neurons, we used multiplexed FISH to quantify the expression of *Cre* alongside the *Sst*, *Peg10*, *Rbfox1*, and *Vgf* transcripts described above. *Cre* expression in the *Sst*-Cre line is observed in 3.43% of DAPI+ LHb cells (Fig. 1A) and was largely confined to the HbX subregion in the dorsal/posterior regions of the habenula (Fig. 1B-C). Approximately 92% of *Cre*+ neurons also expressed *Sst*, either alone (∼70%) or in combination with *Peg10* (15%) and/or *Rbfox1* (8%) (Fig. S2A-H). Additionally, *Sst* expression per cell was positively correlated with *Cre*, while *Peg10* and *Rbfox1* showed an inverse relationship, having low levels of expression in cells that had high levels of *Cre* (Fig. S2C-E). We quantified penetrance of *Cre* expression in each LHb subpopulation by examining the percentage of each subpopulation that expressed *Cre*. We found *Sst+* neurons expressed *Cre* in the largest proportion (16.4%) while *Peg10+* (3.6%) and *Rbfox1+* (1.4%) expressing cells coexpressed *Cre* in low numbers (Fig. S2B)*. Cre*+ cells in the *Kcnc1-*Cre line represented 21.79% of all DAPI+ cells (Fig. 1D) and were distributed throughout the anterior/posterior axis of the LHb with densest expression in the most medial and lateral extents of the LHb, while avoiding the HbX (Fig. 1D-F). Most cells expressing *Cre* within the *Kcnc1*-Cre lin*e* also expressed *Vgf* (23%), *Rbfox1* (19%), or both (26%) (Fig. S2I). A small proportion of *Cre+* cells also expressed *Peg10* (10%), but this was typically in the rare cells that also co-expressed either *Rbfox1, Vgf,* or both (Fig. S2I). Furthermore, *Peg10* expression showed an inverse relationship with *Cre* expression in individual cells (Fig. S2K), while both *Vgf* and *Rbfox1* showed weak positive correlations (Fig. S2L-M). Finally, in approximately 26% of *Cre+* cells we did not detect any of the three transcripts we tested, indicating either expression of our chosen genes was too low in these cells, they are a non-neuronal group, or they represent a subpopulation labeled by genes other than were used here (Fig. S2I). We quantified *Cre* penetrance and found penetrance lowest in *Peg10+* (29.4%) cells, but in a large proportion of *Rbfox1* (51.6%) and *Vgf* (47.8%) cells for *Kcnc1*-Cre mice (Fig. S2J). Within the *Prokr2*-Cre line, *Cre* expression is observed in 7.46% of DAPI+ cells. Similar to the *Kcnc1*-Cre line, *Cre* expression in the *Prokr2*-Cre line was distributed throughout the anterior/posterior axis of the LHb with densest expression in the most medial and lateral extents of the LHb, while avoiding the HbX (Fig. 1G-I). Most cells expressing *Cre* also expressed *Vgf* (38%) or *Rbfox1* and *Vgf* (42%) (Fig. S2Q). A small proportion of *Cre+* cells also expressed *Peg10* (18%), but this was largely seen in cells that also co-expressed either *Rbfox1, Vgf,* or both (Fig. S2Q). Furthermore, *Peg10* expression showed an inverse relationship with *Cre* expression in individual cells (Fig. S2S), while both *Vgf* and *Rbfox1* showed weak positive correlations (Fig. S2T-U). In *Prokr2*-Cre mice, *Cre* penetrance was found in a similar proportion across *Peg10* (18.4%), *Rbfox1* (17.2%), and *Vgf* (15.5%) expressing cells. Together these data demonstrate that transgenic mouse lines can differentially target genetically and spatially distinct populations of neurons in the LHb. The *Sst*-Cre line targets a subpopulation located within the HbX with high specificity but low penetrance. By contrast, the *Cre* expression in the *Kcnc1*-Cre and *Prokr2*-Cre lines avoids the HbX region and instead targets cells expressing *Rbfox1, Vgf,* or dual *Rbfox1*/*Vgf* expressing neurons, with the penetrance of *Cre* expressing in these cell populations varying between each line.

**Figure 1.**
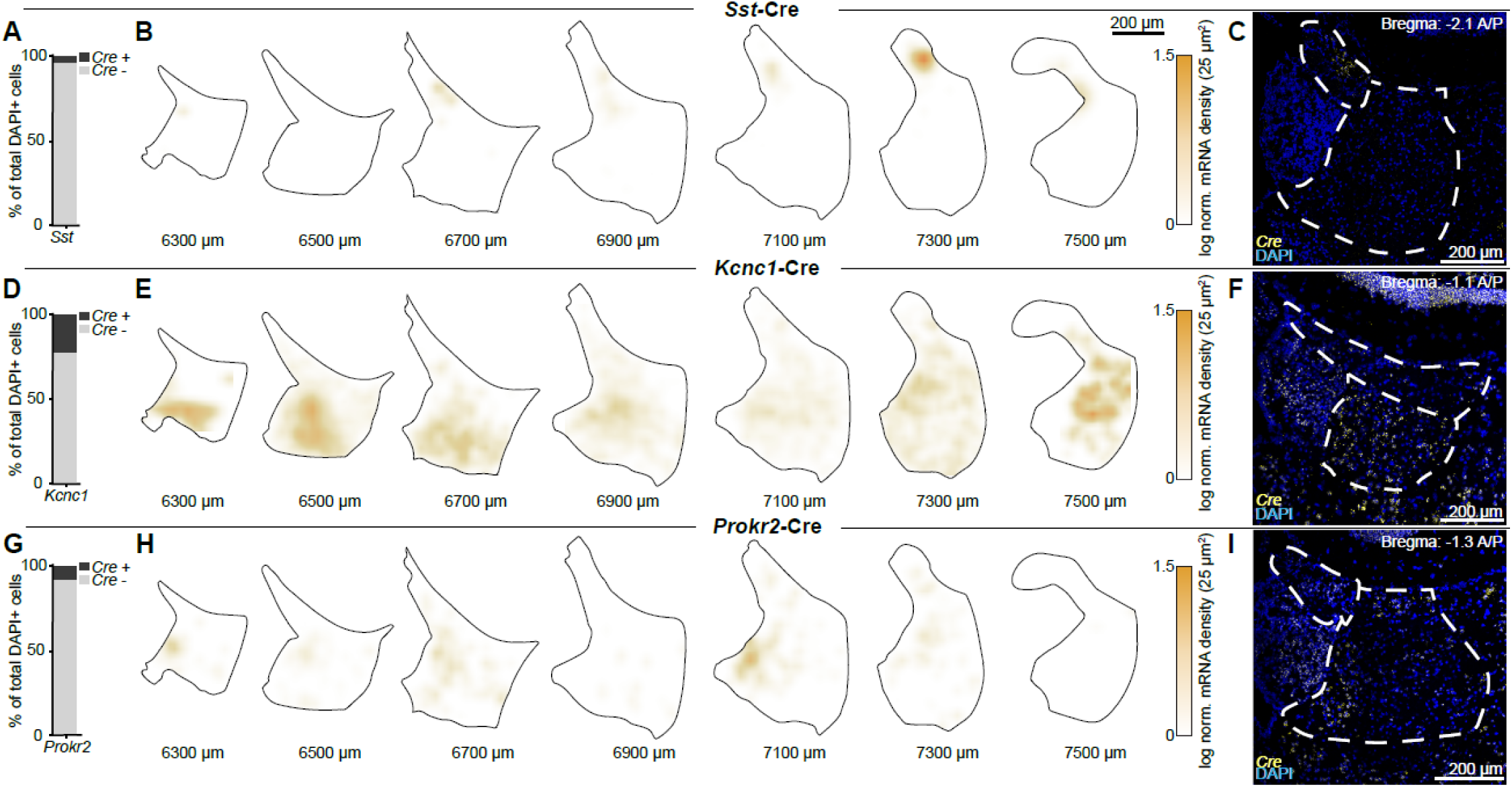
Spatial location and density of *Cre* expression across the LHb differ between transgenic mouse lines. (**A**) Cre-recombinase expressing cells in *Sst*-Cre mice represent 3.43% of all DAPI+ cells (n=29,069 cells; 3 mice) detected within the LHb. (**B**) Heatmap of log normalized *Cre* mRNA density in *Sst*-Cre mice across the stria medullaris, lateral habenula, and habenula commissure in 200µm bins from the CCFv3 rostro-caudal position from 6300µm (approximate bregma AP: -1.0mm) to 7500µm (approximate bregma AP: -2.2mm). (**C**) Representative image of *Cre* mRNA expression in *Sst*-Cre mouse tissue. (**D**) Cre-recombinase expressing cells in *Kcnc1*-Cre mice represent 21.79% of all DAPI+ cells (n=32,753 cells; 3 mice) detected within the LHb, significantly higher (F(2,5)=12.929, p=0.011) than was detected in *Sst*-Cre (p=0.010) and *Prokr2*-Cre mice (p=0.042). © Heatmaps of log normalized *Cre* mRNA density in *Kcnc1*-Cre mice across the rostro-caudal axis. (**F**) Representative image of *Cre* mRNA expression in *Kcnc1*-Cre mouse tissue. (**G**) Cre-recombinase expressing cells in *Prokr2*-Cre mice represent 7.46% of all DAPI+ cells (n=18,093 cells; 2 mice) detected within the LHb. (**H**) Heatmaps of log normalized *Cre* mRNA density in *Prokr2*-Cre mice across the rostro-caudal axis. (**I**) Representative image of *Cre* mRNA expression in *Prokr2*-Cre mouse tissue. Statistics represent a one-way ANOVA with post-hoc Tukey’s HSD.

### Neuron subpopulation-specific electrophysiological properties in the LHb

Differential gene expression analysis from LHb sc-seq datasets suggests significant variability the expression of voltage-gated ion channels across LHb neuronal subpopulations.^32, 33^ Additionally, previous *in vitro* and *in vivo* electrophysiological studies have observed considerable diversity in both passive membrane properties and spontaneous AP firing patterns within the LHb.^22, 40, 41^ We hypothesized that the genetically defined neuronal subpopulations targeted by *Sst*-Cre*, Kcnc1*-Cre *and Prokr2*-Cre mouse lines would also show distinct electrophysiological characteristics. We targeted the LHb with intracranial injections of an adeno-associated virus (AAV) encoding for a Cre-dependent red fluorescent protein (tdTomato; tdTom) to label somata of *Cre*+ neurons in each transgenic mouse line (*Sst*+, *Kcnc1*+, *Prokr2*+; Fig. 2A). Subsequent patch-clamp electrophysiological recordings were obtained from fluorescently labeled neurons across the LHb (Fig. 2B) to measure passive and active membrane properties. Neurons for all mouse lines had approximately similar, and relatively high (>1GΩ), membrane resistance (Fig. 2C). *Sst*+ neurons had a lower membrane capacitance (C_m_) and faster membrane time constant © than both *Kcnc1*+ and *Prokr2*+ neurons (Fig. 2D-E). *Sst*+ neurons also fired fewer APs in response to positive current injections with a greater coefficient of variation (CV) of their ISI than both *Kcnc1*+ and *Prokr2*+ neurons, indicating that the timing of current evoked APs in the *Sst*+ population was less regular (Fig. 2G-I). The AP threshold for all subpopulations targeted was similar, however, the *Sst*+ neurons had slower APs with increased full-width at half height (FWHH) and slower maximum rates of decay on the falling phase of the AP (Fig. 2K-O). Together these measurements suggest that, despite a faster membrane time constant, *Sst*+ neurons fire fewer, slower APs than *Kcnc1*+ and *Prokr2*+ neurons. The passive and active membrane properties of *Kcnc1*+ and *Prokr2*+ neurons were indistinguishable, except *Prokr2+* cells showed reduced attenuation of AP amplitude during positive current injection (Fig. 2H).

**Figure 2.**
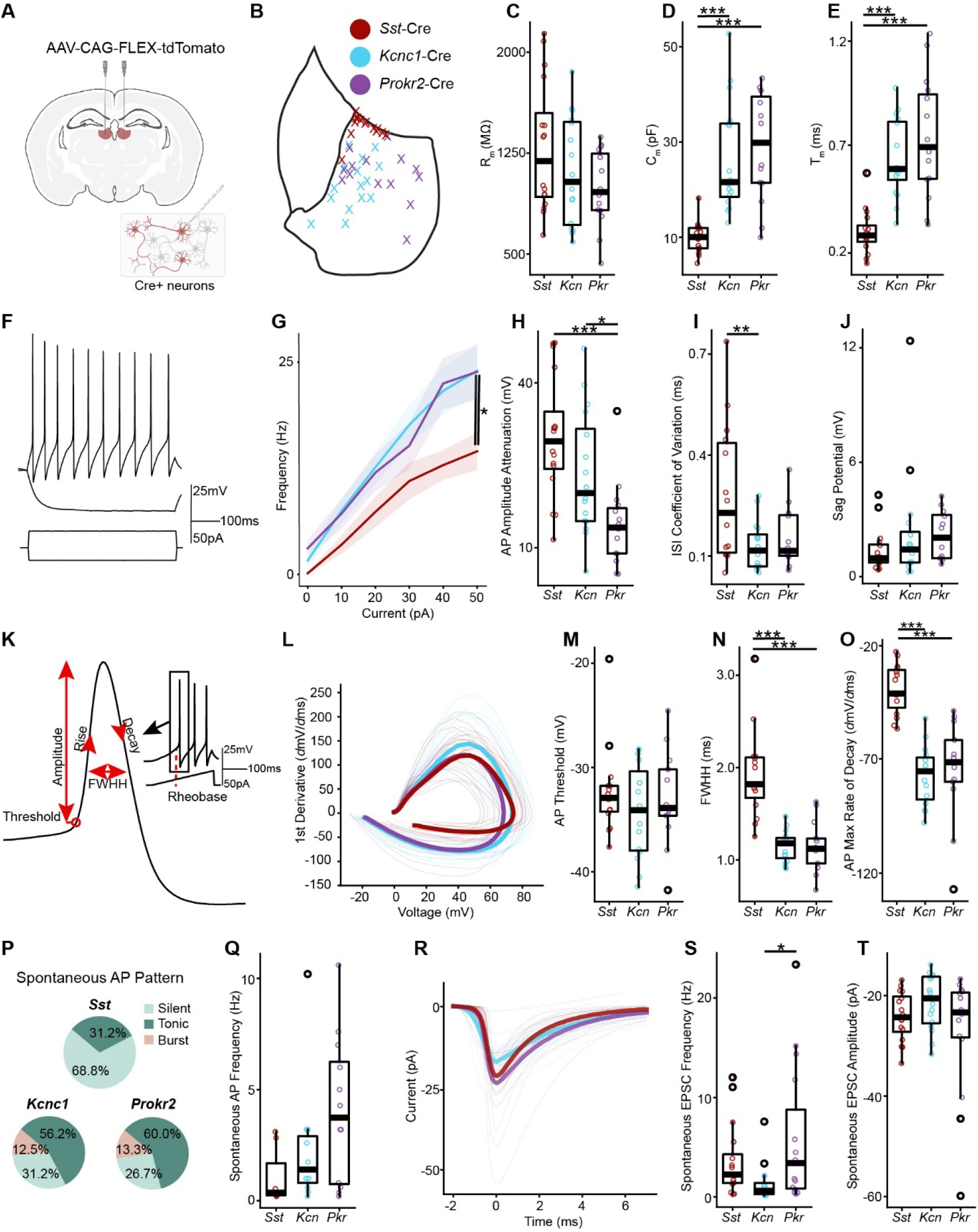
Differential electrophysiological properties across *Sst*+, *Kcnc1*+ and *Prokr2*+ LHb neuronal subpopulations. (**A**) Injection of Cre-dependent AAV encoding tdTomato to target neuronal subpopulations for whole-cell slice electrophysiology recordings. (**B**) Spatial location of patched neurons in Sst (*red*) (n=16 cells), Kcnc1 (*blue*) (n=16 cells), and Prokr2+ (*purple*) (n=15 cells) transgenic mouse lines. (**C**) Membrane resistance (MΩ) was not seen to differ across subpopulations (F(2,44)=2.273, p=0.115). (**D**) Membrane capacitance (pF) was significantly lower (F(2,44)=19.023, p<0.001) in *Sst*+ cells as compared to *Kcnc1*+ and *Prokr2*+ cells, p<0.001 and p<0.001, respectively. © Membrane tau (ms) is significantly lower (F(2,44)=21.338, p<0.001) in *Sst*+ cells as compared to *Kcnc1*+ and *Prokr2*+ cells, p<0.001 and p<0.001, respectively. (**F**) Representative traces of current (*bottom*) and voltage (*top*) recordings during long square current injections. (**G**) Frequency (Hz) of APs during 500ms of positive current injection from 0pA to 50pA differed across subpopulations (F(2,41)=4.391, p=0.019) as *Sst*+ neurons show reduced excitability compared to *Kcnc1*+ and *Prokr2*+ neurons, p=0.038 and p=0.041 respectively. (**H**) *Prokr2*+ neurons had significantly smaller (F(2,44)=9.392, p<0.001) difference in amplitude (mV) between the first and last AP evoked during 500ms of 50pA current injection when compared to *Sst*+ and *Kcnc1*+ neurons, p<0.001 and 0.043 respectively. (**I**) Coefficient of variation (ms) of the ISI across 500ms of 50pA current injection is significantly higher (F(2,39)=5.222, p=0.010) in *Sst*+ as compared to *Kcnc1*+ neurons (p=0.01). (**J**) Sag potential (mV) calculated during -50pA current injection does not differ across subpopulations (F(2,41)=0.959, p=0.392). (**K**) Representative AP plot detailing electrophysiology metrics of interest. (**L**) Phase plot displaying the mean voltage (bold line) by the first derivative of the voltage for the first current-evoked AP at rheobase across all cells. Starting AP voltage was normalized to zero for comparison. (**M**) AP threshold (mV) for current-evoked AP at rheobase did not differ across LHb neuronal subpopulations (F(2,40)=0.778, p=0.466). (**N**) Full-width at half height for current-evoked APs at rheobase was significantly wider (F(2,40)=26.295, p<0.001) for *Sst*+ neurons as compared to *Kcnc1*+ and *Prokr2*+ neurons, p<0.001 and p<0.001 respectively. (**O**) Maximum rate of decay (*d*mV/*d*ms) during current-evoked APs at rheobase is significantly lower (F(2,40)=27.140, p<0.001) in *Sst*+ neurons as compared to *Kcnc1*+ and *Prokr2*+ neurons, p<0.001 and p<0.001 respectively. (**P**) Pattern of spontaneous APs in each neuronal subpopulation across silent, tonic, and burst patterns (Χ^2^(4)=7.59, p=0.108). (**Q**) Frequency (Hz) of spontaneous APs did not differ across subpopulations (F(2,27)=2.727, p=0.083). © Average sEPSC shape across cells in each neuronal subpopulation. (**S**) Frequency (Hz) of sEPSCs recorded at -70mA is significantly lower (F(2,44)=3.767, p=0.031) for *Kcnc1*+ cells as compared to *Prokr2*+ cells (p=0.023). (**T**) Amplitude (pA) of sEPSCs recorded at -70mA did not differ across subpopulations (F(2,44)=2.124, p=0.132). For box and whisker plots, boxes represent the three quartiles (25%, 50%, and 75%) of the data whiskers represent 1.5*IQR and circles are outliers, horizontal line is the median. All statistics represent one-way ANOVAs, except frequency by current comparisons which used two-way repeated measures ANOVA, and post-hoc Tukey’s HSD.

We then examined spontaneous AP firing patterns across the different subpopulations. *Sst*+ neurons had low spontaneous activity with no cells showing a burst pattern and only 31% exhibiting spontaneous tonic firing (Fig. 2P-Q). Conversely, both the *Kcnc1*+ and *Prokr2*+ populations showed three spontaneous AP firing patterns (silent, tonic, and burst) in relatively similar proportions (12-13% burst, 55-60% tonic, and 25-30% silent; Fig. 2P). To confirm if neurons that did not produce spontaneous burst-pattern APs were capable of burst-firing, we injected a hyperpolarizing square wave current (-50pA; Fig. S3A-B). Similar to spontaneous AP firing patterns, no *Sst*+ neuron exhibited rebound burst-firing (Fig. S3B). By contrast, 68% of *Kcnc1*+ neurons produced rebound AP bursts following hyperpolarization, indicating that a larger proportion of *Kcnc1*+ neurons were capable of burst-pattern APs then were suggested by their spontaneous firing patterns. Almost all (93%) of *Prokr2*+ neurons showed rebound burst-firing, indicating that most neurons within this subpopulation can be driven to fire burst-pattern APs (Fig. S3A-B).

Finally, we performed voltage-clamp recordings (V_hold_=-70mV) to examine spontaneous excitatory post-synaptic current (sEPSC) frequency and amplitude to determine if these neuronal subpopulations received differential spontaneous excitatory synaptic input. We did not detect differences in the amplitude of sEPSCs between groups; however, *Prokr2*+ neurons did have an increased frequency of sEPSCs as compared to *Kcnc1*+ neurons (Fig. 2R-T). Together these data support genetic studies showing differential ion channel gene expression between *Sst+* (*Sst-Cre*) and *Vgf+* or *Rbfox1+* (*Kcnc1*-Cre *and Prokr2*-Cre) neuronal subpopulations.^32–34^ Furthermore, they indicate that most *Vgf+* and *Rbfox1+* neurons targeted using the *Kcnc1*-Cre *and Prokr2*-Cre lines are capable of firing APs in a burst pattern.

### Differential axonal targeting of downstream regions by LHb subpopulations

Recent studies have shown that the *Sst*+ subpopulation of LHb neurons sends strong axonal projections to the paranigral nucleus of the VTA^35^. However, if other genetically distinct LHb populations target spatially distinct downstream regions remains unknown. Axonal projections for *Kcnc1+* and *Prokr2+* neuronal subpopulations were quantified using small (50nL) intracranial injections of an AAV encoding a Cre-dependent tdTom and synaptophysin tagged green fluorescent protein (GFP) to label axons and their synaptic terminals, respectively. Slices taken at 100-micron intervals from the LHb to the locus coeruleus were aligned to the Allen Brain Institute common coordinate framework (CCFv3), using the *Aligning Big Brains and Atlases (ABBA)* software. This allowed us to map tdTom positive somata and labeled axons to standardized 3D coordinates and compare their location across animals and mouse lines. Injections into the *Kcnc1*-Cre line showed variations in the density of tdTom positive somata across the medial to lateral axis, with the majority of cells localized to either the lateral or medial fringes of the LHb (Fig. S4A-B). In contrast, injections into the *Prokr2*-Cre line showed tdTom positive somata localized to the lateral edges of the LHb (Fig. S4C-D), with labeled cells shifted laterally and ventrally when compared to those labeled in the *Kcnc1*-Cre (Fig. S4E). Consistent with *Cre* expression observed using FISH (Fig. 1D-H), fewer neurons overall were labeled using the *Prokr2-Cre* line than the *Kcnc1-Cre* line likely labeling a more discrete laterally located subregion of LHb (Fig. S4E).

The pattern of laterally shifted labeling in the *Prokr2*-Cre line was also present when we examined the distribution of axonal labeling (Fig. 3A and 4B). Axons labeled from the *Prokr2*-Cre line overall were shifted more laterally and dorsally than those from the *Kcnc1*-Cre line (Fig. 4B). Axonal projections were quantified as the percent of total structure volume with tdTom positive axons normalized within each animal, or as the percentage of the total detected axons, indicating notable differences in the downstream targets of *Kcnc1*+ and *Prokr2*+ LHb neuronal subpopulations (Fig. 4A). The total detected volume of each downstream target was calculated for each animal and did indicate a slight reduction in the total volume captured for the laterodorsal tegmental nucleus and dorsomedial tegmental nucleus in *Prokr2*-Cre mice (Fig. 4A). This difference did not appear to significantly alter results as these regions did not receive large projections from either subpopulation. Broadly, *Prokr2+* neurons more densely targeted laterally located structures ipsilateral to the injection, including the parabrachial pigmented nucleus and parainterfasicular nucleus of the VTA, while *Kcnc1+* axons were denser medial in the paranigral nucleus of the VTA and the interpeducular nucleus (IPN). Similarly, *Prokr2+* axons were most dense in the dorsal raphe subdivisions (interfascicular, dorsal, lateral, ventral and caudal linear nucleus), while *Kcnc1+* axons were the overwhelming input to the medial and paramedian raphe. While not delineated in the Kim unified anatomical atlas, the RMTg has been identified as a major target of the LHb that is critical for its modulation of VTA dopamine neurons.^13, 42, 43^ This GABAergic structure includes the posterior pole of the VTA immediately lateral to the IPN (corresponding broadly with the parainterfasicular and parabrachial pigmented nuclei of the VTA) and appears heavily innervated by *Prokr2+* and to a lesser extent *Kcnc1+* axons (Fig. 4A, 9.5mm).^13^ Together, the *Kcnc1+* axons generally targeted downstream structures located more medially and ventrally than the *Prokr2+* axons, even when accounting for interindividual differences in the medial-lateral axis location of targeted somata suggesting that LHb neurons labeled with these two lines differentially modulate downstream structures (Fig. S4E-F).

**Figure 3.**
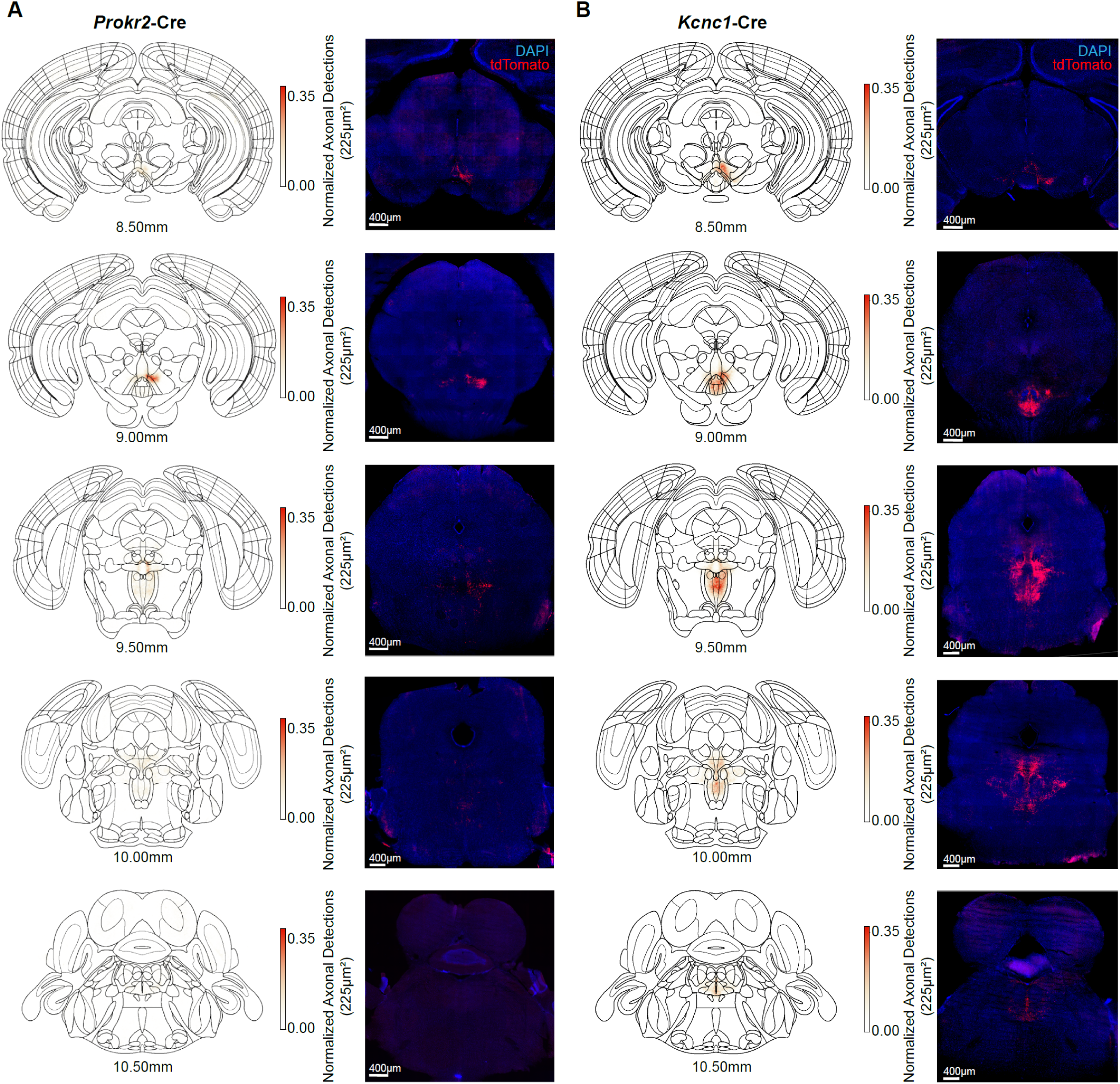
Mapping of axonal densities in downstream structures for *Kcnc1*+ and *Prokr2*+ LHb neurons. (**A**) Heatmaps (*left*) of average normalized axonal detections across all *Prokr2*-Cre mice (n=5) in 500µm segments from CCFv3 rostro-caudal position of 8500µm (approximate bregma AP: -3.3mm) to 10500µm (approximate bregma AP: -5.3mm) and representative images (*right*). (**B**) Heatmap (*left*) of average normalized axonal detections across all *Kcnc1*-Cre mice (n=5) and representative images (*right*).

**Figure 4.**
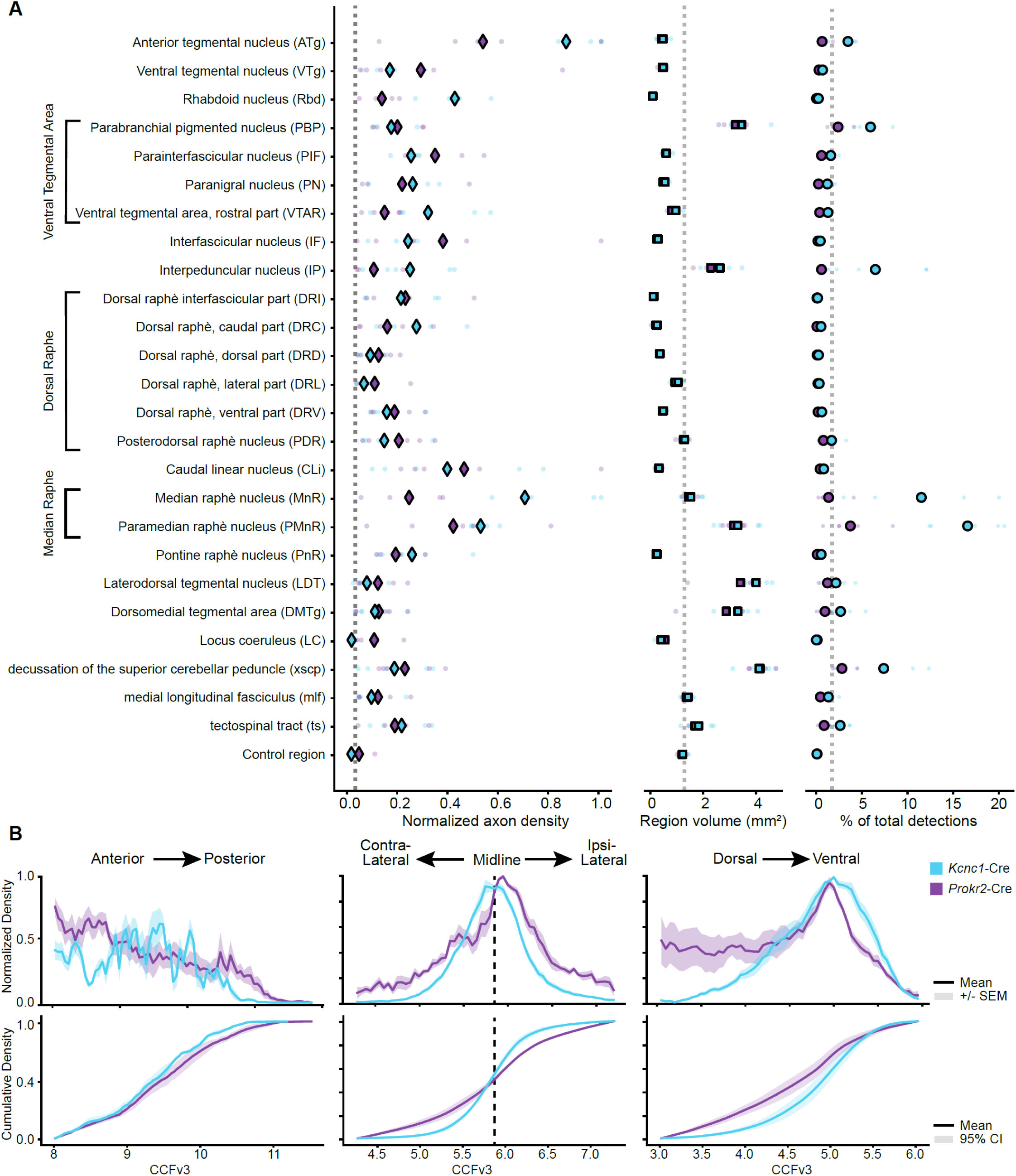
Differential targeting of downstream structures by *Kcnc1*+ and *Prokr2*+ axons. (**A**) Density of axonal detections across downstream regions as measured by the percentage of the structure’s volume positive for tdTom or GFP positive axons, normalized within animal, and then averaged for each transgenic line (Control region: Interstitial nucleus of Cajal) (*left*). The average structure volume utilized in axonal detection analyses (*middle*). Density of LHb projections in each downstream structure as a percentage of total detections (*right*). Larger, dark shaded points indicate the group mean. Lighter points indicate animal means. (**B**) The normalized distribution (*top*) and cumulative distribution (*bottom*) for *Kcnc1*+ and *Prokr2*+ axonal projections across the anterior to posterior (*left*), medial to lateral (*middle*), and dorsal to ventral (*right*) axes. Dashed lines in medial to lateral plots (*middle*) represent the interhemispheric midway point in the CCFv3 framework (5725µm). Solid lines (blue and purple) represent group means while shaded regions indicate the standard error of the mean (SEM; *top*) or the 95% confidence interval (CI; bottom). The interaction between the spatial position across the given axis and the group (i.e. *Kcnc1*-Cre vs *Prokr2*-Cre) explains a significant amount of variance in normalized axonal density in the anterior to posterior (F(26.03)=12.19, p<0.001), medial to lateral (F(21.71)=45.13, p<0.001), and dorsal to ventral (F(28.29)=295.24, p<0.001) axes.

### Behavioral performance of transgenic mice during a probabilistic switching task

Neurons within the LHb are activated by a wide variety of aversive stimuli and predictive cues which, on a population level, constitute a nRPE signal critical for motivated behaviors and reward learning.^15, 44–50^ Many studies, however, have shown additional diversity where individual LHb neurons can be activated by reward, show biphasic responses of inhibition followed by excitation, or show diverse responses to aversive stimuli depending on location within LHb.^44, 47–49, 51^ We predicted that the genetic and spatial diversity of neuronal subpopulations targeted by the *Kcnc1*-Cre and *Prokr2*-Cre lines described above may reflect differential roles for these subpopulations during behavior. To test this hypothesis, we employed a probabilistic switching task (also known as a two-armed bandit task; 2ABT), a reinforcement learning task which has been shown to require reward-related neural circuits, including midbrain dopamine centers such as the VTA.^52, 53^ Water restricted, freely moving animals are placed in a behavioral arena with three nose-poke ports. A center poke initiates a trial, then the animal makes a choice, moving to the left or right-side ports to probabilistically receive a water reward (∼2uL, Fig. 5A), with one side delivering rewards at high probability (p_reward_ = 0.9) and the other at low probability (p_reward_ = 0.1). Following the outcome (reward/no reward), the mouse exits the side port, ending the trial. The next trial begins when the mouse returns to the center port (Fig. 5A). Lights above the ports indicate when the center or side ports are active to assist in training but provide no information regarding the location of the highly rewarded port. After receiving 50 rewards, the reward probabilities for the two side ports switch (block transition, dotted vertical line, Fig. 5B). There is no cue that a transition to a new block has occurred and the time and number of trials it takes each animal to reach 50 rewards varies; therefore, following a block transition, the probability that the animal chooses the highly rewarded port (p(high port)) drops dramatically. A well-trained animal adjusts its choices, and p(high port) gradually increases for the next 10-15 trials (Fig. 5C).

**Figure 5.**
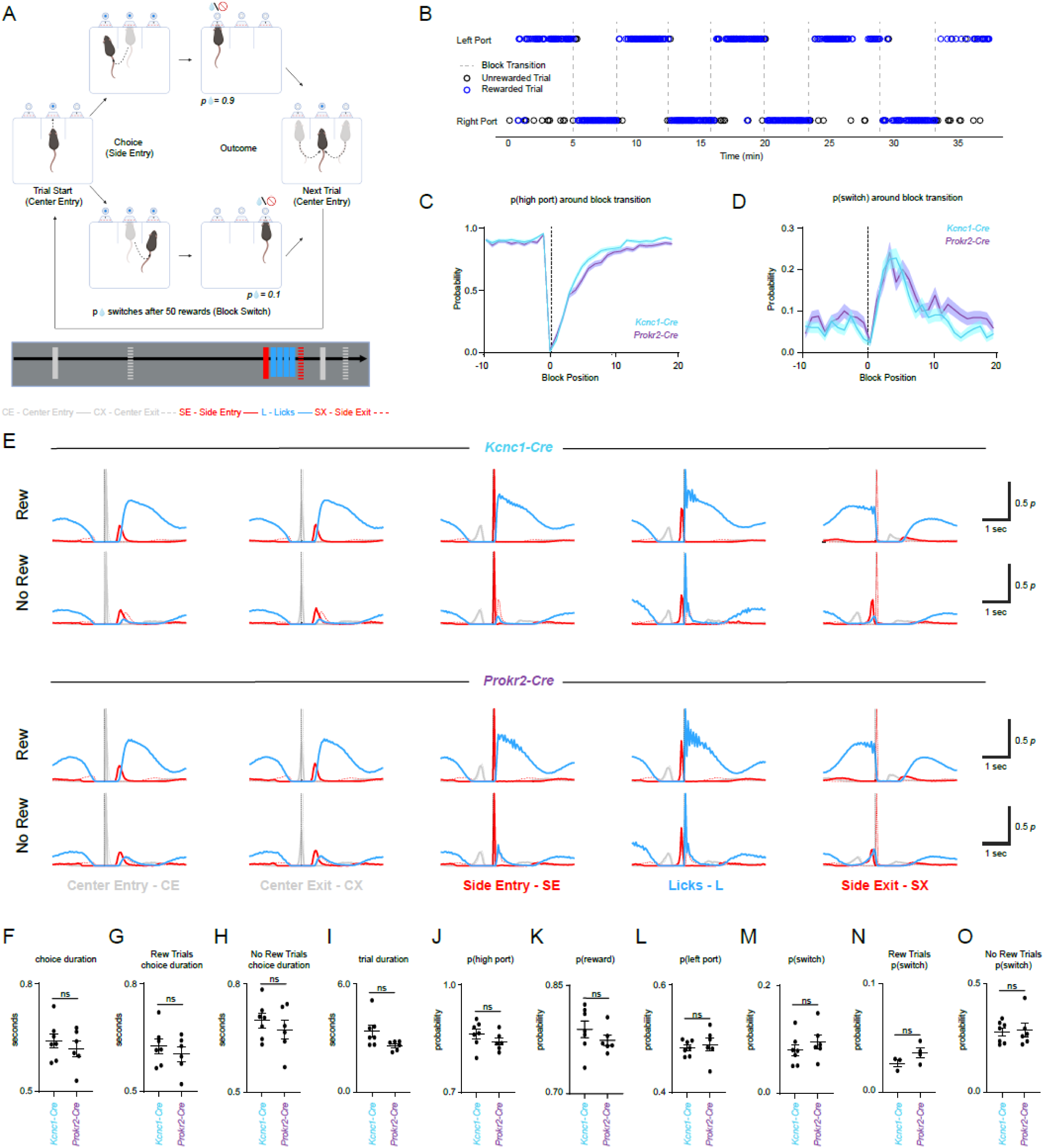
Behavioral performance of *Kcnc1*-Cre and *Prokr2*-Cre mouse lines during a probabilistic switching, a two-armed bandit task (2ABT). (**A**) Diagram outlining performance on the 2ABT. Briefly, a trial begins when a mouse breaks an IR beam in the center nose poke to begin the task. The mouse then chooses a port, left or right, pokes the chosen port, either receives a reward or not then exits the side port ending the trial. The next trial begins when the mouse returns to the center port. After receiving 50 rewards, the probabilities that each port will give a reward (90% and 10%) switch sides (block transition). (**B**) Example of single, 40-minute session of 2ABT performance showing rewarded and unrewarded trials with block transition by a *Prokr2*-Cre mouse. (**C**) Probability of choosing the highly reward port around the block transition. Data are presented as mean (solid line) ± SEM (shaded region). (**D**) Probability of choosing a different port than the one chosen on the previous trial (switching) around the block transition. Data are presented as mean (solid line) ± SEM (shaded region). © 5 second windows (2 seconds prior to center and 3 seconds after) detailing task event probabilities (as probability) across all animals and trials centered at Center Entry, Center Exit, Side Entry, First Lick, and Side Exit (black dotted line) for all reward and unrewarded trials for *Kcnc1*-Cre and *Prokr2*-Cre mice. (**F-O**) Behavioral metrics for *Kcnc1*-Cre and *Prokr2*-Cre mice in the 2ABT. Data are presented as mean ± SEM. Statistical significance is determined via two-tailed nested t-test. (**F**) Choice Duration (Center Exit to Side Entry) for all trials. Ns p=0.4920 (t=0.7108, df=11, F=0.5053). (**G**) Choice Duration following rewarded trials. Ns p=0.4578 (t=0.7695, df=11, F=0.5921). (**H**) Choice Duration following unrewarded trials. Ns p=0.4521 (t=0.7796, df=11, F=0.6078). (**I**) Trial Duration (Center Entry to Center Entry) for all trials. Ns p=0.0550 (t=2.147, df=11, F=4.609). (**J**) Probability of choosing the highly rewarded port for all trials. Ns p=0.2465 (t=01.224, df=11, F=1.498). (**K**) Probability of receiving a reward for all trials. Ns p=0.3045 (t=1.077, df=11, F=1.160). (**L**) Probability of choosing the left port for all trials. Ns p=0.5588 (t=0.6029, df=11, F=0.3635). (**M**) Probability of switching for all trials. Ns p=0.3840 (t=0.9067, df=11, F=0.8222). (**N**) Probability of switching for all rewarded trials. Ns p=0.2290 (t=1.274, df=11, F=1.622). (**O**) Probability of switching for unrewarded trials. Ns p=0.8671 (t=0.1713, df=11, F=0.2933). Behavioral data consists of 5-6 2ABT sessions combined and consists of *Kcnc1*-Cre n=7 (4 female, 3 male) and 21,460 trials and *Prokr2*-Cre n=6 (2 female, 4 male) and 15,463 trials. These behavioral sessions also had photometry recorded simultaneously. Ns = no significance. Diagrams made with Biorender.

We examined baseline performance between *Kcnc1*-Cre and *Prokr2*-Cre mice on the 2ABT. Approaching a block transition, both genotypes perform similarly, choosing the highly rewarded port ∼80% of the time in the ten trials prior to a block transition. At the block transition, both genotypes slowly begin to favor the side with the highly rewarding port as they learn the new port probabilities, adjusting their choices similarly (Fig. 5C). Likewise, the probability that mice of either genotype will switch side ports on subsequent trials (*p(switch)*) is ∼10% in the ten trials leading up to the block transition. P(switch) rises to a similar peak between both genotypes following the block transition and subsequently decays down to ∼10% as the mice adjust to the new block (Fig. 5D). On rewarded and unrewarded trials, both *Kcnc1*-Cre and *Prokr2*-Cre mice show grossly similar behavior for rewarded/unrewarded trials. Following side entry (SE), both *Prokr2-Cre* and *Kcnc1-Cre* mice have similar licking behavior on rewarded and unrewarded trials (Fig. 5E). The time it takes animals to move between the center port exit and side port entrance (choice duration), was similar in *Prokr2*-Cre and *Kcnc1-*Cre mice for all trials regardless of the outcome (rewarded/non-rewarded) of the previous trial (Fig. 5G, H), with trials following rewarded trials having faster choice duration than those following unrewarded trials, as expected based on previous 2ABT studies.^54^

To determine if the genotypes had varying levels of task proficiency, we examined the probability of choosing the highly rewarded port and the probability receiving a reward for both *Kcnc1*-Cre and *Prokr2*-Cre. These metrics showed no significant difference by genotype (Fig. 5J, K). Additionally, the *Kcnc1*-Cre and *Prokr2*-Cre mice did not display a port bias (Fig. 5L), with both groups showing a similar probability of switching ports overall (choosing different ports on subsequent trials) or following rewarded or unrewarded trials (Fig. 5M-O). Overall, the two transgenic mouse lines described here perform similarly on the 2ABT, making it unlikely that changes observed in neural activity in the two lines are due to behavioral differences 2ABT performance.

### Differential LHb subpopulation activity during a probabilistic switching task reflects distinct task components

We injected Cre-dependent AAVs into the LHb, encoding calcium sensor GCaMP8f and static fluorophore tdTom or GCaMP8f alone, to selectively label each neuronal subpopulation.^55^ Following injection, a fiber optic probe was implanted above the injection site (Fig. 6A). Probe placement was over the left hemisphere LHb, defining ipsilateral (ipsi; left) and contralateral (contra; right) ports and movements (Fig. 6C, left diagram). Fiber photometry was performed during multiple 2ABT sessions to measure calcium-dependent fluorescence changes, as a proxy for neuronal activity. Comparing simultaneously recorded stable fluorophore (tdTom) and GcaMP8f signals in shows that, while there are small motion artifacts likely due to fiber optic patch cable movement and/or hemodynamic responses, these artifacts contribute a very small component to the observed calcium signal (Fig. S7A-D).^56^ LHb neuronal dynamics during a single 2ABT session show changes within a trial during port choice and outcome (Fig. 6B). Grouped event-triggered averages were aligned to side entry (SE) to capture the outcome period as well as to center exit (CX) to capture the choice period. A window of 1 second (grey shaded box) following the aligned event was used to calculate Δ Peak. Event triggered averages include a 100ms delay (green shaded box) to account for reward delivery (Fig. 6C, right diagram, SE aligned signals shown). On rewarded trials, we observed a significant difference between LHb neural activity of *Kcnc1*-Cre and *Prokr2*-Cre mice. *Kcnc1+* neuronal activity shows a dip following reward while *Prokr2+* activity slowly rises during reward consumption before returning to baseline (Fig. 6D). On unrewarded trials, both subpopulations increase activity with *Prokr2+* neurons showing a significantly stronger and sharper increase in activity following reward omission (Fig. 6E) while *Kcnc1+* neurons rise more slowly, peak much later and then remain elevated for ∼2 seconds. When comparing rewarded and unrewarded trials, *Kcnc1+* neurons show a significant reduction in activity on rewarded trials and an increase on unrewarded trials (Fig. 6F). Conversely, *Prokr2+* neurons show a sharp, immediate spike in activity following reward omission, while rewarded trials lead to a slow rise in activity that returns to baseline after reward consumption (Fig. 6G). Using long dividers to extend the duration between events did not meaningfully affect neuronal activity centered at side port entrance on rewarded or unrewarded trials for either genotype (Fig. S5A-D).

**Figure 6.**
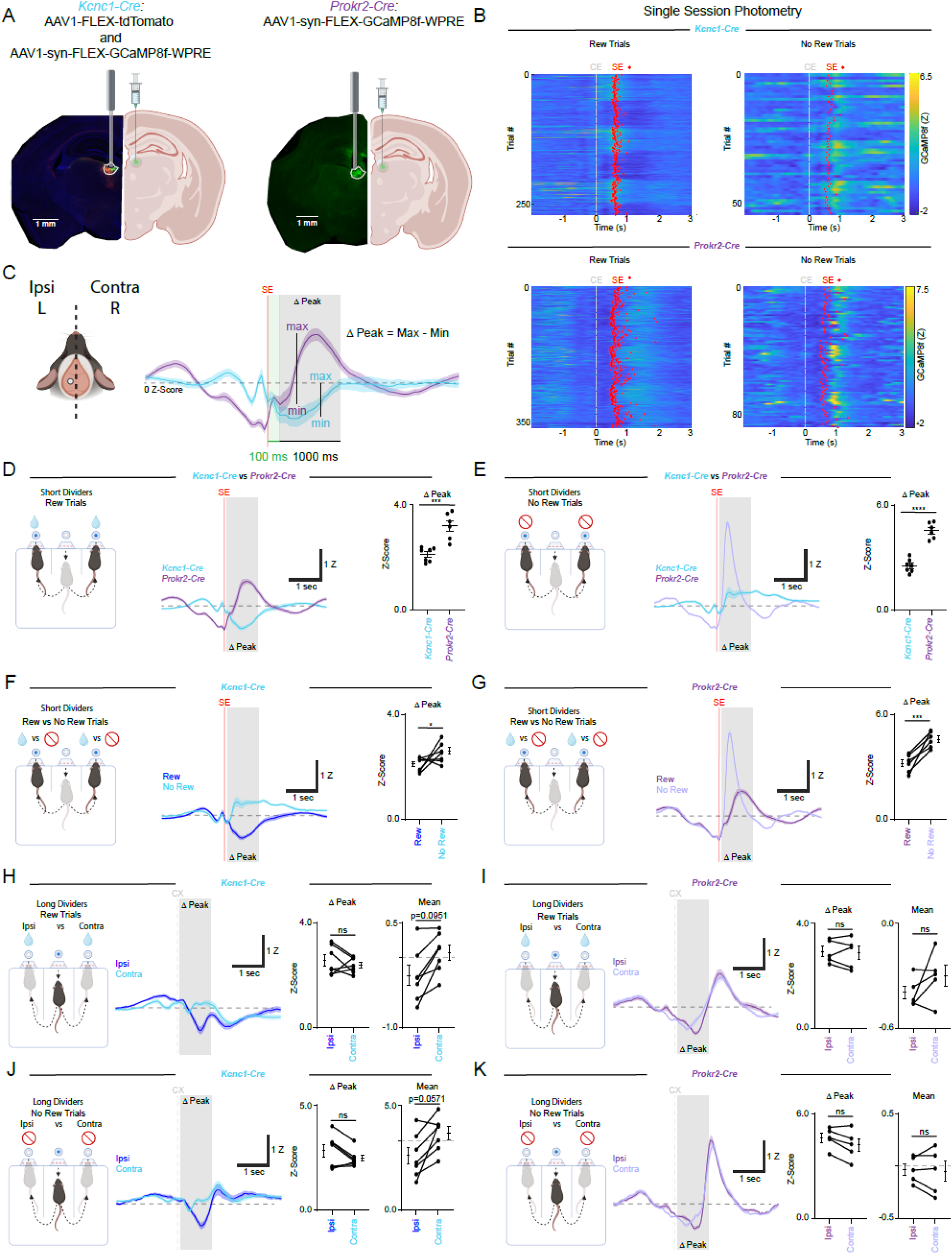
*Kcnc1*+ and *Prokr2*+ neurons show differential activity during directional movement and outcome during a two-armed bandit task (2ABT). (**A**) Example brain slices of *Kcnc1*-Cre mice injected with Cre-dependent AAVs expressing GcaMP8f and tdTom and *Prokr2*-Cre mice injected with Cre-dependent AAV expressing GcaMP8f with probe placement. (**B**) Heatmaps showing a 5 second window (2 before center and 3 after) for a single 2ABT training session’s rewarded and unrewarded photometry by trial for *Kcnc1*-Cre and *Prokr2*-Cre mice centered on Center Exit. Red dots indicate Side Entry. Data are Z-scored with warmer colors indicating larger Z. (**C**) Diagram (left) of example fiber optic probe implant for photometry in the left hemisphere LHb defining ipsilateral and contralateral movements relative to the probe implant. Diagram (right) showing example photometry GcaMP8f traces. Data are presented as Z-score (solid line) ± SEM (shaded region) for a 5 second window (2 seconds before center (vertical red line) and 3 seconds after). Dotted gray line represents zero Z. Green shaded region represents 100 ms after center while gray box represents a 1 second region of data for which Δ Peak is calculated. Δ Peak is given as the maximum value reached in the gray shaded region minus the minimum value. Mean represents the mean photometry activity during the 1 second gray shaded region. (**D – K**) Calcium photometry data are aligned to indicated behavioral event (SE, CX, etc.). For *Kcnc1*-Cre 47 short divider and 41 long divider 2ABT sessions were combined and consists of *Kcnc1*-Cre n=7 (short and long) (4 female, 3 male) animals and for *Prokr2*-Cre 30 short divider and 25 long divider 2ABT sessions were combined across n=6 (short) or 5 (long) (2 female, 4 male for short dividers, 2 female, 3 male for long dividers) animals. For short and long dividers, *Kcnc1*-Cre consisted of 16879 and 10237 trials respectively and *Prokr2-*Cre consisted of 12145 and 9707 trials respectively. For photometry activity traces, data are Z-scored and line graphs presented as described in (**C**). Graphs quantify Δ Peak as mean ± SEM. Statistical significance is determined via two-tailed nested t-test. Diagrams to the left of data indicate trial epoch, dividers used, mouse position, and alignment point of photometry traces. (**D**) Neuronal activity of *Kcnc1*-Cre vs *Prokr2*-Cre mice aligned to Side Entry on rewarded trials using short dividers. *** p=0.0004 (t=4.948, df=11, F=24.48). © Neuronal activity of *Kcnc1*-Cre vs *Prokr2*-Cre mice aligned to Side Entry on unrewarded trials using short dividers. ****p<0.0001 (t=8.719, df=11, F=76.02). (**F, G**) Neuronal activity aligned to Side Entry comparing rewarded to unrewarded trials using short dividers. *Kcnc1*-Cre mice * p=0.0299 (t=2.462, df=12, F=6.063). *Prokr2*-Cre mice *** p=0.0006 (t=4.876, df=10, F=23.78). (**H, I**) Neuronal activity aligned to Center Exit comparing ipsilateral vs contralateral choice for rewarded trials using long dividers. *Kcnc1*-Cre mice Δ Peak ns p=0.4957 (t=0.7026, df=12, F=0.4936), Mean ns p=0.0951 (t=1.812, df=12, F=3.283). *Prokr2*-Cre mice Δ Peak ns p=0.8689 (t=0.1704, df=8, F=0.02904, Mean ns p=0.2139 (t=1.350, df=8, F=1.823). (**J, K**) Neuronal activity aligned to Center Exit comparing ipsilateral vs contralateral choice for rewarded trials using long dividers. *Kcnc1-*Cre mice Δ Peak ns p=0.3068 (t=1.067, df=12, F=1.140), Mean ns p=0.0571 (t=2.105, df=12, F=4.429). *Prokr2*-Cre mice Δ Peak ns p=0.4063 (t=0.8766, df=8, F=0.7684), Mean ns p=0.8625 (t=0.1788, df=8, F=0.03198). ns=no significance. Diagrams made with Biorender.

Aligning photometry signals to center exit (CX) allows for comparison of activity patterns that accompany movements related to choice. For this analysis, long dividers were used to extend the duration between behavioral events providing clearer temporal separation of the individual trial events (Fig. 6H-K, S5E-F). *Kcnc1+* neurons showed dip in activity during movements to the ipsilateral reward port (Fig. 6H-J, S6C). Examining the mean photometry signal during a one second window shows a trend for a difference during ipsilateral vs contralateral choice regardless of trial’s outcome suggesting a movement component to this signal (Fig. 6H and J). Comparatively, *Prokr2+* activity following CX did not differ during directional movements towards the reward port. Examining the mean photometry signal following CX shows no change between side choice for either rewarded or unrewarded outcomes (Fig. 6I and K). We then compared activity during ipsiversive or contraversive movements following CX between *Kcnc1-Cre* and *Prokr2-Cre* mice. We found differences in the Δ Peak following CX between *Kcnc1+* and *Prokr2+* neurons for both ipsilateral and contralateral movements, indicating they show differential activity during movement towards the reward port (Fig. S6A and B). Together, these results suggest additional directional movement related activity in the *Kcnc1+* neurons, while *Prokr2+* neurons do not show differential activity to ipsilateral vs contralateral movements.

To more specifically examine the relationship between movement direction and neural activity, we compared two different epochs when the animal made ipsiversive or contraversive movements on either short (Fig. S6E-H) or long (Fig. S6I-L) dividers. The two epochs defined for ipsiversive movement include one following side exit (SX) of the right port and another following center exit (CX) on trials to the left reward port (Fig. S6E, F, I, and J). In *Kcnc1*-Cre mice using long dividers, dips in fluorescence occurred during ipsiversive movements for both epochs and were not significantly different (Fig. S6I). Contraversive movements in *Kcnc1+* mice also show similar activity patterns regardless of divider size rising after port exit and then dipping as if following movement around a divider (Fig. S6G and K). *Prokr2*-Cre mice on long dividers, with better temporal separation of behavioral events, showed differences in the mean photometry signal as the animal moved towards the reward port. A significant dip in activity occurred as the animal moved towards the reward port that was not present when the animal made a similar directional movement toward the center port, suggesting *Prokr2+* activity differs with reward expectation (Fig. S6J and L). Taken together, these data highlight differential activity patterns between the two LHb subpopulations. Activity within *Kcnc1+* neurons relates to both choice/movement direction and outcome, while *Prokr2+* neurons respond more specifically to trial outcome and reward expectation.

### Trial history dependence of LHb activity

To collect maximize reward collection, animals performing the 2ABT must consider their port choice history and the outcomes of previous trials.^57^ As a result, following rewarded trials, p(switch) is low and animals expect to receive a reward if they make the same choice on the next trial (Fig. 5N-O). If a reward is not delivered on the next trial a reward prediction error (RPE) occurs which has been observed in dopamine neurons in the 2ABT.^52, 53^ To test if RPEs could be observed in LHb subpopulations, we examined how trial history influences LHb neuronal activity across *Kcnc1+* and *Prokr2+* neurons. To simplify trial history, we use a nomenclature in which the capitalization of letters represents the trial outcome, with rewarded trials denoted by a capital letter (A/B) and unrewarded trials denoted by a lower-case letter (a/b). Additionally, the choice to stay vs switch ports between trials is denoted by the letter (either A or B), with the letter A denoting that the mouse selected the same port as the previous trial and the letter B indicates a switch to the opposing port. (Fig. 7A).^57^ Lastly the letter order denotes the trial order with the left most letter representing the previous trial and the right most letter representing the current trial. For example, “Aa” would be used to denote a unrewarded trial that was preceded by a rewarded trial at the same port, while “AB” would indicate a rewarded trial preceded by a rewarded trial at the opposite port (Fig. 7A).

**Figure 7.**
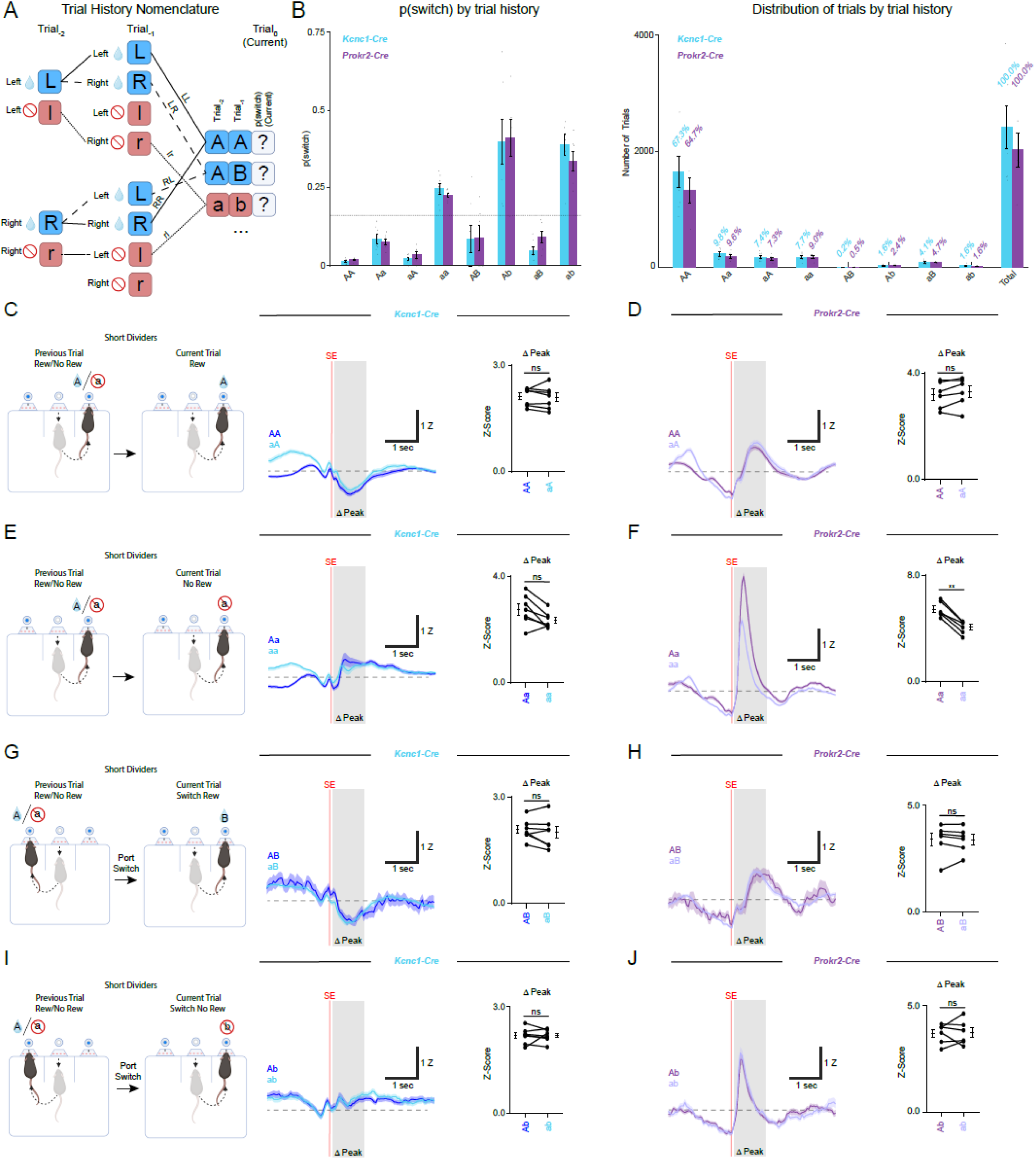
Trial history affects outcome related activity in *Prokr2+*, but not *Kcnc1+* neuronal subpopulations during the 2ABT. (**A**) Diagram outlining nomenclature defining trial history. Capitalized letters indicated rewarded trials. Lower case letters indicate unrewarded trials. Trial history is read right to left with the right most letter indicating the current trial and each letter to the left moving one trial into the past. A’s indicate trials to the same port as the previous trial while B’s are trials that switch port side from the previous trial. R’s are trials to the contralateral port while L’s are to the ispsilateral port. (**B**) Trial history dependence of p(switch) for *Kcnc1*-Cre and *Prokr2*-Cre mice, dashed gray line is mean p(switch) for all trials (*left*). The distribution of different trial histories across all trials (percentages of total trials denoted above each trial type) (*right)*. (**C – J**) Neuronal activity of *Kcnc1-*Cre and *Prokr2*-Cre mice expressing GcaMP8f. Calcium photometry data are aligned to indicated behavioral event (SE, CX, etc.). For *Kcnc1*-Cre 47 short divider divider 2ABT sessions were combined and consists of *Kcnc1*-Cre n=7 (4 female, 3 male) animals and for *Prokr2*-Cre 30 short divider 2ABT sessions were combined across n=6 (2 female, 4 male) animals. *Kcnc1*-Cre mice consisted of 16879 trials and *Prokr2-*Cre mice consisted of 12145 trials. Data are Z-scored and line graphs are presented as mean (solid line) ± SEM (shaded area). Graphs quantifying Δ Peak are presented as mean ± SEM. Statistical significance is determined via two-tailed nested t-test. Diagrams to the left of data show trial type, mouse movement, and center point of line graphs. (**C, D**) Neuronal activity aligned to Side Entry comparing rewarded trials where the previous trial was either rewarded or unrewarded to the same port using short dividers. *Kcnc1*-Cre mice. Ns p=0.8997 (t=0.1288, df=412, F=0.01658). *Prokr2*-Cre mice. Ns p=0.7168 (t=0.3731, df=10, F=0.0.1392). (**E, F**) Neuronal activity aligned to Side Entry comparing unrewarded trials where the previous trial was either rewarded or unrewarded to the same port using short dividers. *Kcnc1*-Cre mice. Ns p=0.1656 (t=1.476, df=12, F=2.180). *Prokr2*-Cre mice. **p=0.0019 (t=4.161, df=10, F=17.32). (**G, H**) Neuronal activity aligned to Side Entry comparing rewarded trials where the previous trial was either rewarded or unrewarded to the opposite port using short dividers. *Kcnc1*-Cre mice. Ns p=0.7042 (t=0.3888, df=12, F=0.1512). *Prokr2*-Cre mice. Ns p=0.9740 (t=0.03339, df=10, F=0.001115). (**I, J**) Neuronal activity aligned to Side Entry comparing unrewarded trials where the previous trial was either rewarded or unrewarded to the opposite port using short dividers. *Kcnc1*-Cre mice. Ns p=0.9514 (t=0.06228, df=12, F=0.002878). *Prokr2-*Cre mice. Ns p=0.8934 (t=0.1374, df=10, F=0.01888). ns= no significance. Diagrams made with Biorender.

Photometry signals were aligned to side port entry (SE) for all analysis on the impact of trial history on outcome related neural activity (Fig. 7C-H). Importantly, p(switch) is strongly modulated by different trial histories indicating that animals consider previous trial outcomes when making their next choice (Fig. 7B). When the current trial is rewarded (AA or aA), trial history has no effect on neural activity in either subpopulation (Fig. 7C, D). When the current trial is unrewarded (Aa or aa), however, there is a significant effect of trial history on activity within *Prokr2+* neurons only. When the previous trial is rewarded (Aa), the increase in activity associated with a reward omission is significantly larger than when the previous trial is not rewarded (aa; Fig. 7E, F). This indicates that expectation differentially impacts neuronal activity and suggests *Prokr2+* neurons display a negative RPE (nRPE) signal. Surprisingly, the nRPE signal is not observed during switch trials, regardless of trial outcome (AB/Ab/aB/ab; Fig. 7G-J). This indicates that expectation may reset when a mouse chooses to switch ports between trials during more exploratory states.

To determine the effects of movement/choice direction on reward history, we modified the trial history nomenclature to preserve port laterality. In this modified nomenclature, instead of A and B, R is used to denote trials to the right (contra) port while L denotes trials to the left (ipsi) port. Again, trial history affects p(switch) strongly, but not differently depending on port chosen (Fig. S8A). During unrewarded trials, neither the selected port (R/L) nor the reward history impacted observed *Kcnc1+* activity (Fig. S8B and D). By contrast, reward history significantly impacted *Prokr2+* activity with unrewarded trials that were preceded by a rewarded trial showing a larger Δ Peak as compared to those preceded by an unrewarded trial (Fig. S8C and E). As described above, there is no effect of trial history during switch trials for either port choice or subpopulation (Fig. S8F-I).

### Generalized linear model of LHb subpopulation activity

Animal movements during a 2ABT trial are complex and occur in quick succession, making it difficult to disambiguate which behavioral events may be associated with specific features of the simultaneously recorded neural signal. Therefore, we trained a generalized linear model (GLM) on the behavioral and photometry data collected during long divider sessions for both *Kcnc1-Cre* and *Prokr2-Cre* animals.^53, 54^ The timing of behavioral events (CS, CX, SE, SX, etc) are used by the model to fit photometry signals. For each behavioral variable, the GLM assigns a kernel of time shifted beta (β) coefficients that represent the contribution of that variable to the photometry signal (GcaMP8f fluorescence; Fig. 8A and C). These kernels can then be convolved with the actual timing of behavior events in a trial and summed to create a “reconstructed” GcaMP8f signal which is compared to the actual (original) signal (Fig. 8A, right). We selected 80% of the data to train the GLM and 20% to test the model. The 80/20 split was randomly assigned across the entire data set ten times (10-fold cross validation) to ensure there was no bias in the model for any one mouse and/or training day to skew the results (Fig. 8B).

**Figure 8.**
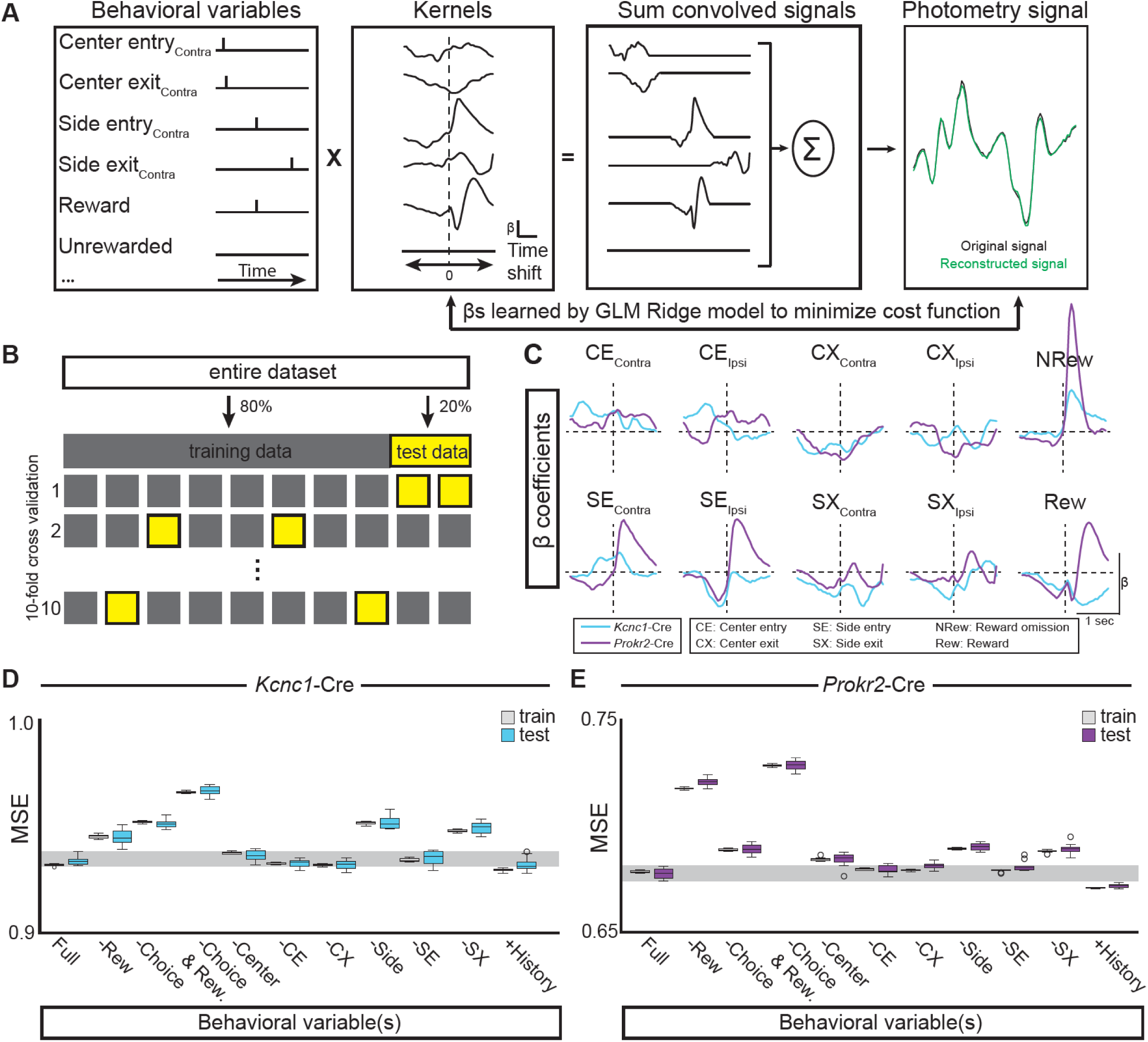
Generalized linear model demonstrates differential contributions of choice, reward, and trial history to activity of LHb subpopulations during behavior. (A) GLM workflow: behavioral variables are convolved with their kernels. Each time shift in the kernel consists of an independent β coefficient fit jointly by minimizing a cost function. The convolved signals are then summed to generate a reconstructed signal which can be directly compared to the original photometry trace. (B) The entire dataset is divided into training and test datasets. The GLM is fit on the training data and evaluated on the test data using mean squared error (MSE). Following a grid search that compared multiple regularization types (ridge, elastic net, ordinary least squared) in combination with a large hyperparameter space, ridge regression (α=1) was found to give the smallest error following cross-validation. (C) Kernels for the behavioral variables included as features in the *Full* GLM. Behavioral predictors gave information regarding choice (Ipsi/Contra), reward, and port entry/exit (blue=*Kcnc1-*Cre, purple=*Prokr2-*Cre). (D, E) Box plots showing MSE for the full model (Full) and models in which the indicated behavioral predictor(s) were omitted (-) or added (e.g. +History) for both the train (gray) and test (blue/purple) datasets (Boxes represent the middle 50% of the data, horizontal lines is the median, and whiskers are 1.5*IQR, outliers are shown as circles, see Materials and Methods section for details). Each model-run uses a different combination of data used for train/test split as illustrated in B.

The β coefficients generated for *Kcnc1+* and *Prokr2+* neural activity showed changes to reward, direction of chosen port (ipsi vs contra), and exit from the side port, reflecting our previous analysis of event-aligned subpopulation activity (Fig. 8C). While the mean square error (MSE) for *Kcnc1+* activity (Fig. 8D) was larger than *Prokr2+* activity, the range of the error was similar between groups (Fig. 8D-E). To determine the contribution of each behavioral event(s), behavioral variables were removed from the training sets and the MSE of the reduced model was compared to that of the full model (termed *Full;* Fig. 8D-E). Based on the increase in MSE compared to the full model, Choice and Side Exit strongly contribute to *Kcnc1+* model fits, reflecting the impact of directional movements on neural activity of this subpopulation (Fig. 8D). By contrast, removing Reward variables more strongly increased MSE in *Prokr2+* model fits (Fig. 8E). The addition of trial history (Fig. 7A-B) designations to reward variables significantly reduced MSE for *Prokr2+* model fit but did not impact MSE for *Kcnc1+* models (Fig. 8D, E). Together, these analyses support the conclusion that *Kcnc1+* activity is more strongly affected by choice/direction of behavioral events, while *Prokr2+* activity is strongly driven by trial outcome and trial history.

## Discussion

Here we link genetically, anatomically, and electrophysiologically defined neuronal subpopulations in the LHb to differential activity patterns during motivated behavior. We propose that genetically distinct LHb subpopulations can be differentially targeted using transgenic mouse lines, but that only neurons in the HbX region (*Sst*-Cre*+*) are distinguishable by their intrinsic electrophysiological properties. Further, neuronal subtypes in the LHb labeled using *Kcnc1-*Cre target medially located downstream structures such as the MRN, while LHb neurons labeled using *Prokr2*-Cre project axons to the VTA and RMTg. Finally, only *Prokr2*+ LHb neurons show evidence to negative reward prediction error responses during a probabilistic switching task, while *Kcnc1*+ LHb neurons are modulated by choice direction and value. We propose that by targeting genetically distinguishable LHb subpopulations we can begin to unravel the diverse functions of the LHb in motivated behavior.

### Genetic categorization of LHb neuronal subpopulations targeted by transgenic mouse lines

Single-cell transcriptomics (sc-seq) studies have described genetically distinguishable neuronal subpopulations in the LHb.^32–34^ Considerably less is known regarding the spatial distribution of these subpopulations particularly along the anterior/posterior axis of the LHb, however several recent studies have highlighted the need to incorporate spatial organization into studies of LHb function. We used four genes (*Sst, Peg10, Rbfox1*, and *Vgf*), which we and others have shown to label genetically distinct subclasses using sc-seq, and mapped their spatial distributions across the entire extent of the LHb as well as their co-expression with *Cre* in three transgenic mouse lines. *Cre* expression patterns in *Sst*-Cre mice largely mirrored the pattern observed by *Sst* expression and can be used to target the HbX population in future studies.^35^ *Cre* expression in *Kcnc1*-Cre and *Prokr2*-Cre mice largely avoided both the central region of the LHb as well as HbX, and subsequently showed minimal co-expression with *Peg10* and *Sst*, which is highly expressed in these regions. Interestingly, *Cre* expression in the *Prokr2*-Cre line shows more overlap with *Vgf* than the *Kcnc1*-Cre line (80% vs 47%, respectively) which is seemly at odds with anatomical experiments showing viral labeling biased towards the lateral portion of the LHb (Fig. S4). However, many of the *Cre*+/*Vgf+* neurons in the *Prokr2*-Cre line also express *Rbfox1* and these triple-labeled cells are likely to be largely restricted to the rostral pole of the LHb and not labeled with our viral injections/coordinates (see Fig. S4C). Together these results have caused us to revisit conclusions regarding labeling of distinct neuronal subclasses with *Vgf* and *Rbfox1* transcripts (Fig. S1B and C). These transcripts show the greatest degree of co-expression in neurons in the rostral pole of the LHb, which have largely evaded further study given the difficulty in performing viral injections and electrophysiological recordings in this area due to its small size and dense myelination. The caudal two thirds of the LHb show less co-expression of *Vgf* and *Rbfox1* and more bias towards the medial or lateral fringes of the LHb, respectively.

### Electrophysiological diversity within the LHb

Of the mouse lines examined here, *Sst*-Cre mice appear the most distinct in their gene expression, spatial location, and electrophysiological properties. These neurons have a comparatively smaller membrane capacitance, slower rate of AP decay, wider APs, and do not appear to possess the capacity to fire AP bursts spontaneously or in response to hyperpolarizing current injection. Differences in AP shape are likely driven by differential expression of sodium (*Scn8a*) and potassium (*Kcnc1-4, Kcna2*) channels, reported in previous sc-seq results. ^32, 33^ Additionally, while sc-seq does not indicate expression of genes related to AP bursting (*Cacna1g-©, Hcn1-4*, and *Grin1)* within *Sst+* neurons, additional research is needed to determine if the expression of these genes is modulated by experience or acute/chronic stress. Other studies indicate these cells project to the paranigral nucleus of the VTA, increase in activity in response to rewards, and contribute to reductions in anhedonic behavior in mice.^35^ The difference between activity observed in this *Sst* subpopulation and that of the majority of reports from LHb is representative of the underlying heterogeneity that has been incompletely captured to date.

The *Kcnc1*-Cre and *Prokr2*-Cre mouse lines had yet to be explored as potential tools for targeting LHb neuronal subpopulations. Though neurons within these subpopulations display similar active and passive membrane characteristics, they differ in their probability of burst-pattern AP following hyperpolarizing current injection, with the vast majority of *Prokr2+* neurons producing single or continuous AP bursts in response to hyperpolarization. As changes in the rates of burst-pattern APs within the LHb both *in vitro* and *in vivo* are consistently observed following chronic stress paradigms and appear causally related to anhedonia in mice, *Kcnc1*-Cre and *Prokr2*-Cre mice may provide a more targeted approach to the neuronal populations most directly impacted.^16^

### Projection targets and subcircuits of the LHb

We observed projections from neurons in the *Prokr2*-Cre line targeting the VTA (subregions include PBP, PN, PIF and adjacent IF) as well as strong projections to the PMnR which collectively encompass regions that include RMTg. While not labeled in mouse brain atlases, the RMTg location has been outlined in prior neuro-anatomical experiments.^13, 42^ Conversely, projections from the *Kcnc1*-Cre line densely innervate several subregions of the MRN (MnR, PMnR, PnR, Atg) and caudal DR (DRC). Overall, this results in axonal patterns shifted medially for the *Kcnc1*-Cre line and laterally for the *Prokr2*-Cre line. Anatomically these patterns suggest at least two parallel circuits arising from the LHb, the laterally-located “canonical” circuit that innervates the GABAergic RMTg to modify VTA dopamine neuron firing,^58^ while the other medially-located circuit primarily innervates serotonergic centers.^59, 60^ The existence of other parallel circuits is likely, as we did not examine the centrally located *Peg10+* neurons of the LHb, these neurons also express *Gpr151* and *Htr2c* and may project more broadly to downstream regions.^32, 61, 62^ Future studies will focus on examining specific synaptic connections between these distinct presynaptic populations and downstream dopamine, serotonin, and GABA neurons.

### Activity patterns of LHb neurons reveal distinct relationships to motivated behavior

During probabilistic switching tasks, like the 2ABT, mice must integrate information regarding choice outcomes and update ongoing strategies to successfully identify the more highly rewarded port. Neurons within the LHb are activated by a wide variety of aversive stimuli and predictive cues including foot shock,^44, 45^ quinine,^44^ social aggression/defeat,^44, 46^ looming stimuli,^47^ air puffs,^48^ and reward omission.^15, 49, 50^ However, studies also show additional diversity where individual LHb neurons can be activated by reward, show biphasic responses of inhibition followed by excitation, or vary their responses depending on medial/lateral location within LHb.^44, 47–49, 51^ We hypothesized that the diversity of LHb activity patterns observed during rewarding and aversive stimuli may be explained by genetically and spatially defined neuronal subtypes targetable with transgenic mouse lines. We found that *Prokr2+* neurons showed strong, rapid, and consistent increases in activity in response to reward omission. Interestingly, these neurons also showed increases in activity to reward, however these responses were smaller in magnitude and showed slower dynamics. By examining reward history dependent effects, we also found a strong influence of an animal’s reward outcome history on the magnitude of the reward omission response, indicating that these neurons are specifically sensitive to outcomes that are worse than expected. As these neurons send dense projections to the RMTg and VTA, we hypothesize that this subpopulation contributes to nRPE responses (but not positive) in dopaminergic neurons of the VTA though connectivity with both local GABAergic neurons in VTA and RMTg.^3, 15, 60, 63^ Conversely, *Kcnc1+* neurons showed suppression of activity following a rewarded outcome during the 2ABT which was significantly different than the slow rise observed in the *Prokr2+* neurons. Furthermore, *Kcnc1+* neurons showed a phasic dip in activity as the animal engaged in a contraversive movement, relative to the implanted optic fiber, regardless of whether the motion was towards a side or center port. Finally, responses of *Kcnc1+* neurons to the outcome of a trial were not strongly modulated by trial history, indicating a stricter value-based signal, rather than prediction error (Fig. 8D). Genetically defined activity patterns in LHb subpopulations described here could underlie proposed control of value-based decision-making tasks by providing cellular substrates for choice, value, and prediction error to different downstream targets.^50^

Collectively, these findings link genetic, electrophysiologic, anatomic, and behavioral domains to detail subtype-specific properties within the LHb. Future studies are warranted to investigate how each of these subtypes are modified both during and following chronic stress providing valuable insights into the contribution of specific neuronal subpopulations to the presentation of anhedonic and despair-related behaviors.^23, 46, 64^

## Resource Availability

### Lead Contact

Requests for further information, resources, and reagents should be directed to the lead contact, Michael L. Wallace

### Materials Availability

All reagents generated in this study are available from the corresponding author upon request and completion of a material transfer agreement. Any additional information required to reanalyze the data reported in this work is available from the lead contact upon request.

### Data and code availability

All data supporting the findings of this study are available from the corresponding author upon request.

## Acknowledgments

We thank E. Kraft and A. Natesan for assistance validating FISH probes and protocols, S. Spytek for tissue processing and immunostaining, and J.P. Roussarie for advice and reagents and the Wallace lab for support and critical reading of the manuscript. This work was supported by funding from the NIH (R01-MH133608, R00-NS105883), the Brain Behavior Research Foundation, and the Whitehall Foundation.

## Author Contributions

MC: validation, methodology, investigation, software, formal analysis, visualization, writing – original draft

LH: validation, investigation

JGP: investigation

SS: investigation

JMM: validation, methodology, investigation, software, formal analysis, visualization, conceptualization, writing – original draft

MW: funding acquisition, conceptualization, methodology, supervision, writing – original draft

## STAR Methods

### Key resources table

| Product |  | Manufacturer |
|---|---|---|
| **Mouse Lines** |  |  |
| *Kcnc1*-Cre | (Tg(Kcnc1-cre)MN77Gsat) | MGI:5311735 |
| *Prokr2*-Cre | (B6.FVB(Cg)-Tg(Prokr2-cre)MG17Gsat/Mmucd) | MMRRC_037342-UCD |
| *Sst*-Cre | (B6N.Cg-Ssttm2.1(cre)Zjh/J) | IMSR_JAX:018973 |
| Ai14 | (B6.Cg-Gt(ROSA)26Sortm14(CAG-tdTomato)Hze/J) | IMSR_JAX:007914 |
| C57BL/6J |  | IMSR_JAX:000664 |
| **Viruses** |  |  |
| AAV2/1-phSyn1(S)-FLEX-tdTomato-T2A-SypEGFP-WPRE |  | Addgene #51509-AAV1 |
| AAV2/8-phSyn1-FLEX-tdTomato-T2A-SypEGFP-WPRE |  | Addgene #51509-AAV8 |
| AAV2/8-CAG-FLEX-tdTomato |  | Addgene #28306-AAV8 |
| AAV-syn-FLEX-GCaMP8f-WPRE |  | Addgene #162379-AAV1 |
| **Antibodies** |  |  |
| Chicken anti-GFP |  | Invitrogen A10262 |
| Rabbit anti-mCherry |  | Abcam Ab167453 |
| Goat anti-Chicken Alexa647 |  | Invitrogen A21449 |
| Goat anti-Rabbit IgG Alexa750 |  | Invitrogen A21039 |
| **Fluorescent In situ Hybridization** |  |  |
| ImmEdge® Hydrophobic Barrier Pen |  | Bio-techne 310018 |
| RNAscope™ Multiplex Fluorescent Detection Kit v2 |  | Bio-techne 323100 |
| RNAscope™ Probe - CRE |  | Bio-techne 312281 |
| RNAscope™ Probe- Mm-Rbfox1 |  | Bio-techne 519911 |
| RNAscope™ Probe- Mm-Peg10 |  | Bio-techne 512921 |
| RNAscope™ Probe- Mm-Sst |  | Bio-techne 404631 |
| RNAscope™ Probe- Mm-Vgf |  | Bio-techne 517421 |
| Opal 520 Reagent Pack |  | FP1487001KT |
| Opal 570 Reagent Pack |  | FP1488001KT |
| Opal 690 Reagent Pack |  | FP1497001KT |
| Opal Polaris 780 Reagent Pack |  | FP1501001KT |

## Materials and Methods

### Animals

Mice were housed within the AAALAC accredited BUSM Animal Science Center and were kept on a 12:12hr light cycle. Food and water were provided *ad libitum* unless otherwise stipulated. All procedures were reviewed and approved by the BUSM Institutional Animal Care and Use Committee (IACUC). Transgenic Cre-recombinase mouse lines are from a C57/B6J background and were genotyped at p21 to determine Cre-recombinase expression. Ai14 litters were obtained from homozygous breeders and paired with *Sst*-Cre homozygous males for *Sst*-Cre x Ai14 double transgenic animals used for whole-cell patch-clamp slice electrophysiology experiments described below.

### Fluorescence *in situ* hybridization (FISH)

FISH experiments were conducted using eight mice, 3 *Kcnc1*-Cre (2 male and 1 female), 2 *Prokr2*-Cre (1 male and 1 female), and 3 *Sst*-Cre (2 male and 1 female) that were between p71 and p163 at the time of tissue collection. Mice were anesthetized using isoflurane and euthanized via decapitation. Tissue was collected and frozen at -80° C in Tissue-Tek OCT Compound. Frozen tissue was sliced at 10-micron thickness at -20° C via cryostat and immediately mounted. Mounted sections were maintained at -80° C until staining. FISH was performed using the ACD RNAscope™ Multiplex Fluorescent V2

### Assay and Akoya Biosciences Opal Fluorescent reagents

*Kcnc1*-Cre and *Prokr2*-Cre tissue was stained against *Cre*, *Vgf*, *Rbfox1*, and *Peg10*, while Sst-Cre tissue was stained against *Cre*, *Sst*, *Rbfox1*, and *Peg10*. Mountant (Prolong Gold Antifade) was allowed to cure for a minimum of 72 hours before imaging. Imaging was completed at 4x and 20x via an Olympus VS200 slide scanner, and at 40x on an Olympus FV3000 confocal microscope. 20x and 40x images were registered to the Kim Unified Atlas via the Aligning Big Brains and Atlases (ABBA) plugin on FIJI. Cellular and sub-cellular detections were performed in QuPath. Data processing and figure generation were performed with Python 3.12. Differences in co-expression between *Cre* and additional mRNA transcripts (*Vgf*, *Rbfox1*, and *Peg10*) between *Kcnc1*-Cre and *Prokr2*-Cre animals was determined with a Pearson’s chi-squared test for independence and post-hoc adjusted Pearson residuals with Bonferroni correction for multiple comparison.

### Stereotaxic Surgeries

#### Viral injections

All stereotaxic viral injections were performed following p55 and were conducted under general anesthesia via inhaled isoflurane, 4-5% for induction and 1-2% for maintenance, with sterile technique. For slice electrophysiology experiments, 200nL of adeno-associated viruses (AAVs) expressing Cre-recombinase dependent tdTom (Addgene #51509, titer: 1.63E12 GC/mL, or Addgene #28306, titer: 1.7E12 GC/mL) were injected bilaterally across the lateral habenula (LHb: AP= -1.55mm, ML= +/-0.45-0.48mm, DV= -2.85mm). For axonal tracing experiments, 50nL of AAVs expressing Cre-recombinase dependent tdTom and synaptophysin bound EGFP (Addgene #51509, titer: 1.63E12 GC/mL) were injected unilaterally within the LHb of 6 *Kcnc1*-Cre (1 male and 5 female) and 7 *Prokr2*-Cre (4 male and 3 female). Animals were only included within later tracing analyses if tdTom positive labeled neurons were observed in the LHb. Importantly, we were unable to infect the *Sst-*Cre line with several AAV serotypes (AAV8, AAV9, AAV PHP.eB, or AAVretro) this prevented us from examining both axonal projections and *in vivo* activity for this subpopulation. We suspect this small population of cells may not express the receptor for these viral serotypes or lies too close to the third ventricle to be consistently targeted by viral injection. All viruses were diluted within 0.9% sterile saline. Pulled glass capillaries used for injection were moved to 0.1mm ventral to the injection site for 10 seconds, kept at the target coordinates for 3 minutes before injection, 3 minutes after injection, and finally moved to 0.1mm dorsal to the injection site for 3 minutes before removal. Simple interrupted sutures were used to close the incision site, and mice were kept on a heating pad until they had recovered from anesthesia. Buprenorphine and meloxicam were given subcutaneously for post-surgical analgesia.

#### Fiber photometry

For all fiber photometry experiments the same procedures were used as described above for the viral injection of 200nL of GCaMP8f (Addgene #162379, titer: 1.2E12 GC/mL) and tdTom (Addgene #28306, titer: 4.25E11 GC/mL) in 3 *Kcnc1-*Cre (1 male and 2 female) and 4 *Prokr2-*Cre (2 male and 2 female) mice. Mice were between p75 and p90 for all fiber photometry surgeries. Following viral injection, fiber implants were lowered 0.1mm dorsal to the viral injection (Injection: AP= -1.5mm, ML= -0.4mm, DV = -2.85mm. Probe: AP= -1.5mm, ML= -0.4mm, DV = 2.75mm). Implants were secured to the skull with ethyl cyanoacrylate gel and C&B Metabond quick adhesive dental cement. Mice were allowed to recover from fiber implantation surgery for one week prior to the start of water restriction and behavioral training. Fiber photometry recordings were not performed until at least 3 weeks after surgery to allow sufficient time for viral expression. The frequency of GCaMP8f recordings varied across mice due to technical and physical limitations; however, recordings occurred at three days in a row at maximum to reduce the impact of photobleaching on recorded signal.

### Slice Electrophysiology

Electrophysiology was performed at least three weeks after AAV injection to allow for adequate viral expression. A subset of recordings were performed with *Sst*-Cre x Ai14 mice that were p50 or older at the time of tissue collection. The total number of mice used for electrophysiology experiments includes, 11 *Kcnc1-*Cre mice (5 male and 6 female), 15 *Prokr2-*Cre mice (6 male and 9 female), 3 *Sst*-Cre (2 male and 1 female), mice and 6 *Sst*-Cre x Ai14 mice (4 male and 2 female). Mice were anesthetized using isoflurane before transcardial perfusion with ice cold artificial cerebrospinal fluid (aCSF, 125mM sodium chloride, 2.5mM potassium chloride, 1.25mM sodium phosphate monobasic monohydrate, 25mM sodium bicarbonate, 25mM dextrose, 2mM calcium chloride, and 1mM magnesium chloride). Brain tissue was sliced coronally at 250 micrometer thickness in ice cold aCSF on a vibrating microtome and hemisected before being transferred into a warmed choline recovery solution (92mM choline chloride, 2.5mM potassium chloride, 1.2mM sodium phosphate monobasic monohydrate, 30mM sodium bicarbonate, 20mM HEPES, 25mM dextrose, 5mM sodium ascorbate, 2mM thiourea, 3mM sodium pyruvate, 7mM tris base, 0.5mM calcium chloride, and 10mM magnesium chloride) and warmed aCSF for 10 minutes each. Tissue was allowed to come to room temperature within aCSF for at least 45 minutes before beginning recordings. All recordings were performed in tdTomato positive neurons at room temperature within aCSF, and with a potassium methanesulfonate internal solution (135mM potassium methanesulfonate, 3mM potassium chloride, 10mM HEPES, 4mM magnesium-ATP, 0.3 sodium-GTP, 8mM sodium phosphocreatine, 1mM EGTA, and 1mM calcium chloride). Voltage clamp recordings were conducted at -70 mV. Recordings were not adjusted for junction potential and series resistance compensation was not used. All solutions were continuously oxygenated with 95% O_2_ / 5% CO_2_. Data was acquired with the pCLAMP (Molecular Devices) and initially analyzed with Clampfit (Molecular Devices). Additional analysis and visualization were done in Python 3.12.

### Immunohistochemistry (IHC)

Mice were anesthetized using isoflurane before transcardial perfusion with 1x PBS followed by 4% PFA. Tissue was placed into 4% PFA overnight and then transferred to a 30% sucrose solution for at minimum 48 hours. All tissue was sliced at 50 micrometer thickness coronally on a microtome. Slices at 100-micron intervals were stained with chicken anti-GFP (Invitrogen A10262), rabbit anti-mCherry (Abcam Ab167453), Goat anti-chicken Alexa647 (Invitrogen A21449), and Goat anti-rabbit Alexa750 (Invitrogen A21039) before being mounted with Prolong Diamond Antifade Mountant (Invitrogen P36961). Slices were imaged at 4x and 20x via an Olympus VS200 slide scanner and registered to the Kim Unified Atlas with the ABBA plugin. Cells positive for tdTomato were identified manually in QuPath and downstream axonal density was determined in QuPath via a pixel classifier trained on Cy5 and Cy7 channel intensities. Resulting data was processed and visualized using Python 3.12. Generalized additive models, in R, were used to determine the impact of transgenic line on somata location within the LHb as well as downstream axonal projection targets. Terms including the spatial location had a smoothing basis function applied to account for the non-linear relationship between location and cell or axonal density.

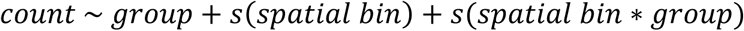

The added variance explained by the location by group interaction term was determined by comparing this model to a reduced GAM, excluding this interaction term, via a one-way ANOVA.

### Probabilistic Switching Task (aka 2-Armed Bandit Task)

The apparatus used for the behavior is as described previously with the following modifications.^54^ Clear acrylic barriers 3 cm (short dividers) or 5.5 cm (long dividers) in length were installed in between the center and side ports to extend the trial time to aid in better behaviorally resolved photometry recordings. Water was delivered in ∼2 μL increments. Hardware and software to control the behavior box is available online: https://github.com/HMS-RIC/TwoArmedBandit and https://edspace.american.edu/openbehavior/project/2abt/.

Following viral injection and fiber implant surgery (see above), mice were water restricted between 0.8 and 1.2 ml per day prior to training and maintained at >80% initial body weight for the full duration of training and photometry. All training sessions were conducted in the dark under red light conditions. During the task a blue LED above the center port signals to the mouse to initiate a trial by poking in the center port. Blue LEDs above the side ports are then activated, signaling the mouse to poke in the left or right-side port within 2 seconds. Side port reward probabilities are defined by custom software (MATLAB) and ranged from 10%-90% depending on the experiment. Withdrawal from the side port ends the trial. The 1 second intertrial interval (ITI) before the center port is active again to start the next trial occurs upon center port entry, insuring that a trial is at minimum 1 second total. An expert mouse can perform 300-600 trials in a 40 min session.

To train the mice to proficiency, animals were subjected to incremental training stages. Each training session lasts for ∼40 minutes, adjusted according to the mouse’s performance. Mice progress to the next stage once they were able to complete at least 100 successful trials with at least a 75% reward rate. On the first day, they were habituated to the behavior box, with water being delivered from both side ports and triggered only by a side port poke. Side ports are baited to encourage side port exploration In the next stage, mice learned the trial structure – only a poke in center port followed by a side port poke delivers water, In this stage, the animal had 15 seconds to select a side port after a center poke started a trial before the center port reset. The center port is baited with a drop of water to encourage center port exploration. Then, the mice transitioned to learning the block structure, in which 30 rewarded trials on one side port triggers the reward probabilities to switch (block transition). In this stage, the reward probabilities are deterministic and the initial center to side interval was 12 seconds. As animals complete >100 successful trials, the window is gradually lowered until they complete >100 trials with a two second window. Once this level of performance is achieved, we began probabilistic reward delivery (p_High_=90%, p_Low_=10%). Mice are then run at this stage until they are well-trained.^54^ For all 2ABT training and experiments, mice performed trials in the presence of barriers in between the center and side ports. *Prokr2*-Cre mice used in this study were also used to help optimize photometry recording procedures, box settings, light power settings, and pharmacology and therefore were more experienced with the 2ABT task than *Kcnc1*-Cre mice.

### Fiber photometry

Fiber implants (Doric, MFC_200/230-0.48_4-5.5mm_MF1.25_FLT) on the mice were connected to a 0.48 NA patchcord (Doric Lenses, MFP_200/220/900-0.48_3-4m_FCM-MF1.25, low autofluorescence epoxy), attached to a filter cube (FMC5_E1(465-480)_F1(500-540) _E2(555-570)_F2(580-680)_S, Doric Lenses). Excitation light from LEDs (Thorlabs) and was amplitude modulated at 167 Hz (470 nm excitation light, M470F3, Thorlabs; LED driver LEDD1B, Thorlabs) and 223 Hz (565 nm excitation light, M565F3, Thorlabs, LED driver LEDD1B, Thorlabs). The following excitation light power measured at the end of the patch cord were used: 470nm=30μW, 565nm=13.5μW for dual fluorophore mice and 470nm=30µW for single fluorophore mice. Signals from the photodetectors were amplified in DC mode with Newport photodetectors (NPM_2151_FOA_FC) and received by a Labjack (T7) DAC streaming at 2000 samples/sec. The DAC also received synchronous information about behavior events logged from the Arduino which controls the behavior box. The following events were recorded: center port entry and exit, side port entry and exit, lick onset and offset, solenoid activation (water delivery), and LED light onset and offset.

### Photometry Analysis

The frequency modulated signals were detrended using a rolling Z-score with a time window of 1 minute (12000 samples). As the ligand-dependent changes in fluorescence measured *in vivo* are small (few %) and the frequency modulation is large (∼100%), the variance in the frequency modulated signal is largely ligand independent. In addition, the trial structure is rapid with an average inter-trial interval of < 3 sec. Thus, Z-scoring on a large time window eliminates photobleaching without affecting signal. Detrended, frequency modulated signals were frequency demodulated by calculating a spectrogram with 1 Hz steps centered on the signal carrier frequency using the MATLAB ‘spectrogram’ function. The spectrogram was calculated in windows of 216 samples with 108 sample overlap, corresponding to a final sampling period of 54 ms. The demodulated signal was calculated as the power averaged across an 8 Hz frequency band centered on the carrier frequency. No additional low-pass filtering was used beyond that introduced by the spectrogram windowing. For quantification of fluorescence transients as Z-scores, the demodulated signal was passed through an additional rolling Z-score (1 min window). To synchronize photometry recordings with behavior data, center port entry timestamps from the Arduino were aligned with the digital data stream indicating times of center-port entries. Based on this alignment, all other port and lick timings were aligned and used to calculate the trial-type averaged data shown in all figures. The Z-scored fluorescence signals were averaged across trials, sessions, and mice with no additional data normalization. Statistical comparisons were made by measuring the mean Z-scored fluorescence signal across a 1s window (Δ Peak) 100 ms after a given behavioral event (CE, SE, SX, …) for all trials per mouse. To aid in photometry and behavior alignment, the first ten seconds of photometry were excluded. Matlab and python analysis scripts were written with the aid of Claude Code.

Statistical analysis of behavioral and photometry data: Outputs from matlab and Jupyter notebook analyses calculating Δ Peak for photometry traces (described in Fig. 6C) or individual value for behavioral data (Fig. 5F-O) were entered into GraphPad Prism 10 in a nested manner with the value for each individual mouse for each trial of a given type or trial epoch entered. An unpaired nested t-test was then performed, taking into account the intra-trial variance in signal/performance for individual mice and the inter-mouse variance within each group.

### Generalized linear model

Photometry recordings and behavioral data used for the GLM analysis (Fig. 8) were collected from *Kcnc1-*Cre (n=7) and *Prokr2-*Cre (n=5) mice as indicated with 5 sessions per mouse and ∼ 400-500 trials/session. These data were aligned to behavioral events to create a predictive matrix *X* (of dimensions *N x F*) and a response vector, **y** (of dimension *N*), where *N* is the number of samples recorded in a session and *F* is the number of behavioral “predictors” in the analysis. The predictors consisted of values 0 and 1 to indicate if a behavioral event (for example a center port entry) occurred in the time bin.

Behavioral predictors listed in Fig. 8D-E are defined as “Rew”=reward delivery or omission, “Choice”=direction (Ipsi/Contra) of side entry, Center=Center entry and exit, CE=Center entry, CX=Center exit, Side=Side entry/exit, SE=Side entry, SX=Side exit, History=reward delivery/omission divided into 8 possible action/outcome combinations including the current trial and one trial in the past (see Fig. 7B for trial history subtypes).

For each predictive matrix, a design matrix *φ*(*X*) (of dimensions *N* × *F* (2*T* + 1)) was constructed from *T* time shifts forward and backward (*T* = 20, 54 ms each) for each feature, allowing the GLM to fit coefficients that corresponded to time-based kernels for each of the predictive features in *X*. Data from the ITI period, in which there are no task-relevant behavioral events, were excluded, and only data spanning shortly before center entry and after side-port exit were modelled. When initial and final time shifts spanned the boundary between two trials, the overlapped data were included twice (once in each of the trials on either side of the boundary) to ensure sufficient representation of each event in training and test datasets.

To optimize the hyperparameters and evaluate the GLMs performance, we performed a grid search across elastic net, ordinary least squares, and ridge regressions. For each model run, a 10-fold group shuffle split (GSS) by trial was applied to the training set to obtain cross-validated ranges for the MSEs, based on an 80– 20 training/test split within each of the 10 GSS folds. Ridge regression (α=1) was determined to be the best performing model, based on the lowest and least variable MSE score (*Kcnc1*-Cre= 0.93, SD=0.002; *Prokr2*-Cre= 0.68, SD=0.002). We then tested the effect of omitting behavioral variables on the GLM performance (Fig. 8D-E) and re-fit the GLM with 10-fold GSS to obtain cross-validated ranges for the MSE values used in the box plots. For the chosen model (Ridge Regression), the algorithm minimizes an associated cost function with respect to the fitted coefficients as follows, where ***J*** is the cost function to be minimized, ***X*** is the design matrix (set of time-shifted behavioral events), ***y*** is the response vector (GCaMP8f), ***β*** is the set of fitted coefficients, **||**a**||^2^_2_**is the sum of the squared entries in vector a, and ***α*** is the regularization parameter.

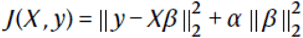

Additional details regarding GLM runs are available in Jupyter Notebooks online at: https://github.com/mwall2017/sabatini-glm-workflow

## Supplemental Figures

**Supplemental Figure 1.**
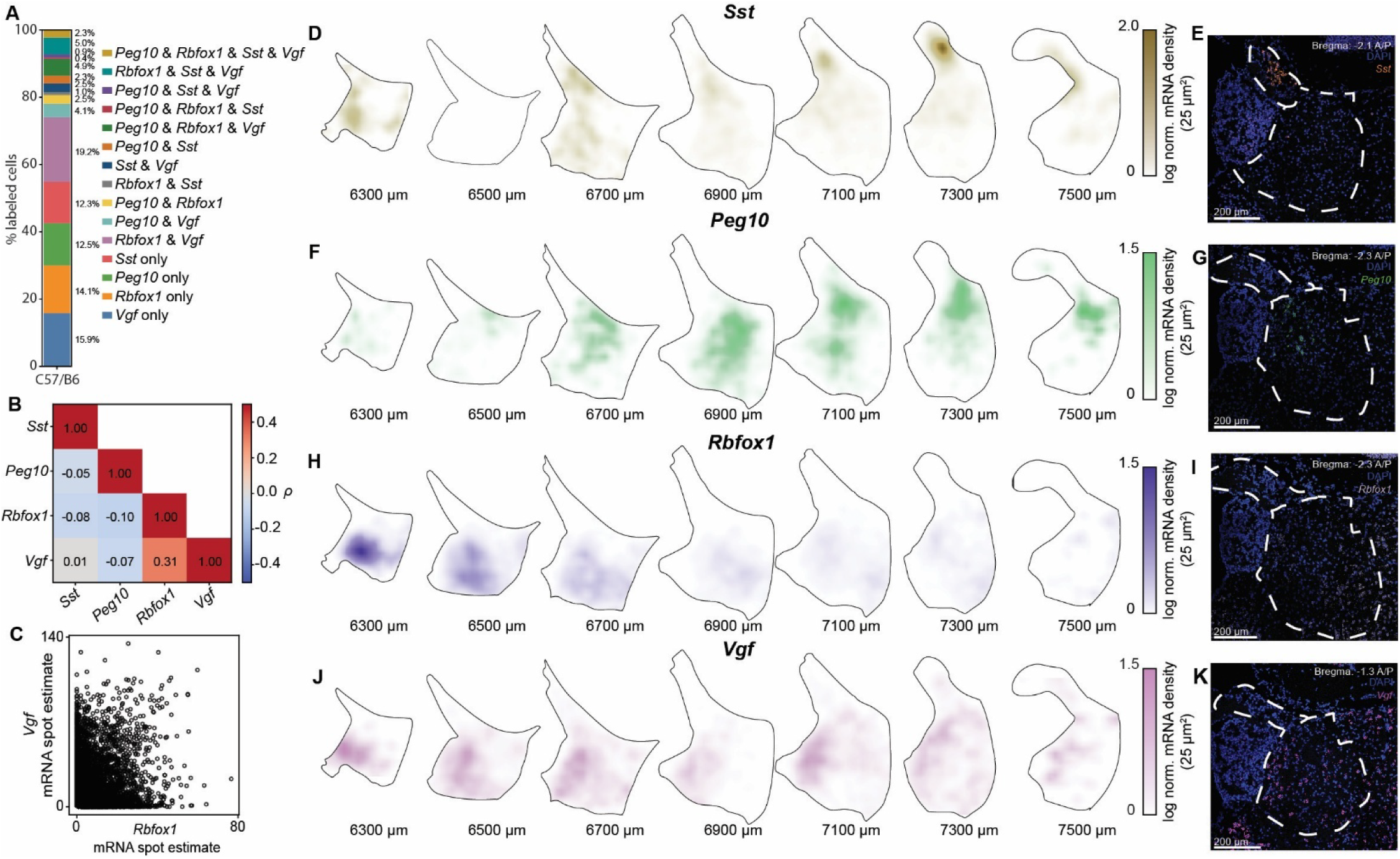
Spatial distribution and cellular expression of transcripts used to mark LHb neuronal subtypes. (**A**) Co-expression of detected mRNA transcripts across labeled cells within wild-type mice (n=11,120 cells; 3 mice). Cells labeled with mRNA transcripts represent 56.7% of all DAPI+ cells. (**B**) Correlation of mRNA expression within cells. Values listed are the relevant Spearman rank correlation rho. (**C**) Scatterplot of mRNA spot estimates for *Rbfox1* and *Vgf* wherein each spot represents one cell. (**D**) Heatmaps of log normalized *Sst* mRNA density across the stria medullaris, lateral habenula, and habenula commissure in 200µm sections from the CCFv3 rostro-caudal position of 6300µm (approximate bregma AP: -1.0mm) to 7500µm (approximate bregma AP: -2.2mm). (**E**) Representative image at 20x of *Sst* mRNA expression at estimated Bregma A/P: -2.1mm. (**F**) Heatmaps of log normalized *Peg10* mRNA density. (**G**) Representative image at 20x of *Peg10* mRNA expression at estimated Bregma A/P: -2.3mm. (**H**) Heatmaps of log normalized *Rbfox1* mRNA density. (**I**) Representative image at 20x of *Rbfox1* mRNA expression at estimated Bregma A/P: -2.3mm. (**J**) Heatmap of log normalized *Vgf* mRNA density across the stria medularis, lateral habenula, and habenula commissure. (**K**) Representative image at 20x of *Vgf* mRNA expression at estimated Bregma A/P: -1.3mm.

**Supplemental Figure 2.**
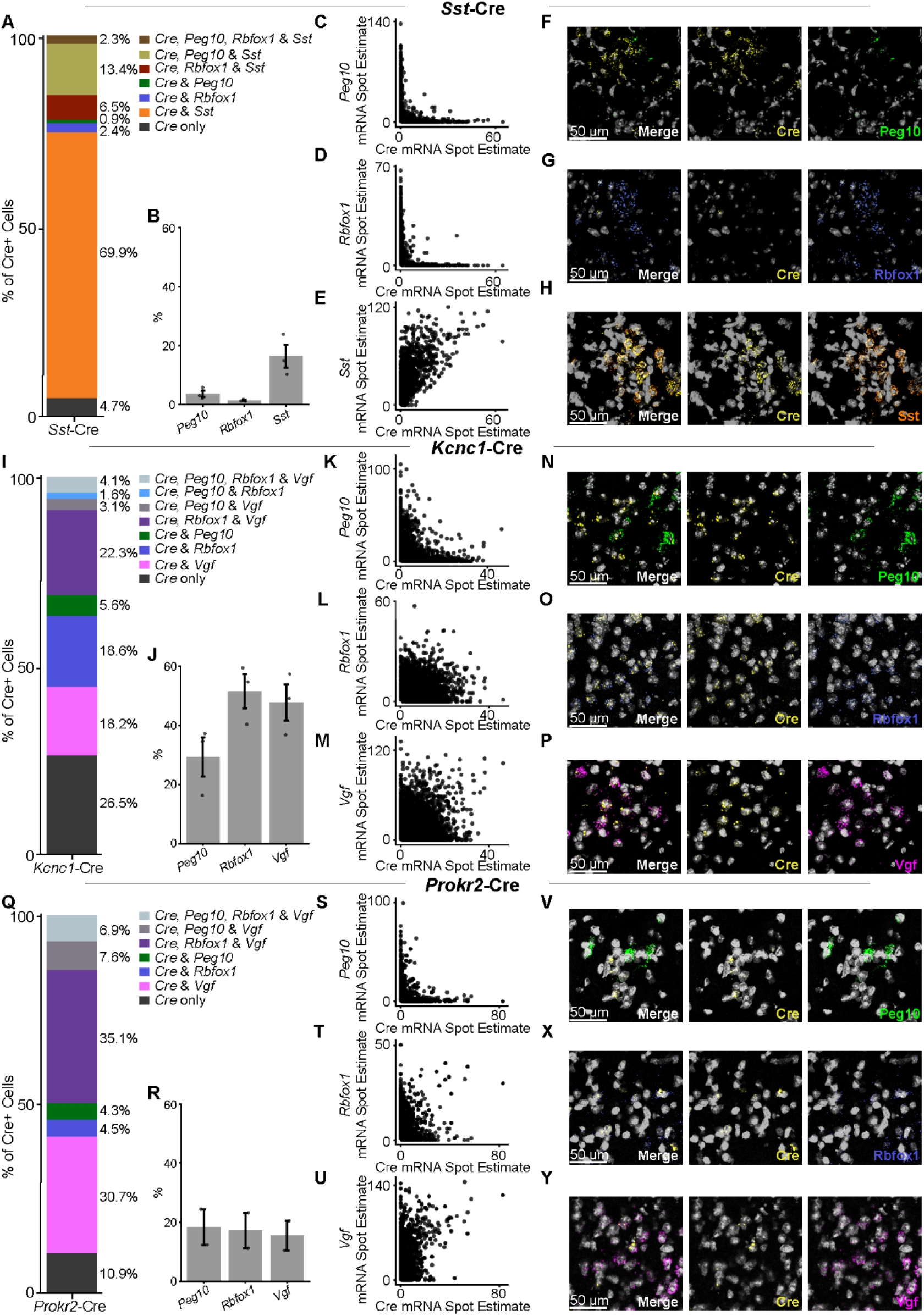
Co-expression of Cre-recombinase and LHb subpopulation transcripts. (**A**) Co-expression of *Cre* and LHb subpopulation mRNA transcripts across *Cre*+ cells in *Sst*-Cre mice (n=980 cells; 3 mice). (**B**) Percent of cells co-expressing *Cre* and each mRNA transcript for *Sst*-Cre mice as a function of all cells positive for each mRNA transcript. (**C**) Relationship between *Cre* and *Peg10* mRNA estimates (n=10,022 cells) in *Sst*-Cre mice. (**D**) Relationship between *Cre* and *Rbfox1* mRNA estimates (n=21,467 cells) in *Sst*-Cre mice. (**E**) Relationship between *Cre* and *Sst* mRNA estimates (n=8,983 cells) in *Sst*-Cre mice. (**F-H**) Representative images at 40x displaying mRNA overlap of *Peg10* (*top*), *Rbfox1* (*middle*), and *Sst* (*bottom*) with *Cre* in *Sst*-Cre mice. (**I**) Co-expression of *Cre* and LHb subpopulation mRNA transcripts across *Cre*+ cells in *Kcnc1*-Cre mice (n=7303 cells; 3 mice). This pattern showed an association with group (Χ^2^(7)=151.69, p<0.001), as *Kcnc1*-Cre mice displaying increased prevalence of *Cre* and *Rbfox1* co-expression (p=0.031) than expected. (**J**) Percent of cells co-expressing *Cre* and each mRNA transcript for *Kcnc1*-Cre mice as a function of all cells positive for that mRNA transcript. (**K**) Relationship between *Cre* and *Peg10* mRNA estimates (n=14,049 cells) in *Kcnc1*-Cre mice. (**L**) Relationship between *Cre* and *Rbfox1* mRNA estimates (n=14,965 cells) in *Kcnc1*-Cre mice. (**M**) Relationship between *Cre* and *Vgf* mRNA estimates (n=15,528 cells) in *Kcnc1*-Cre mice. (**N-P**) Representative images at 40x displaying mRNA overlap of *Peg10* (*top*), *Rbfox1* (*middle*), and *Vgf* (*bottom*) with *Cre* in *Kcnc1*-Cre mice. (**Q**) Co-expression of Cre and LHb subpopulation mRNA transcripts across *Cre*+ cells in *Prokr2*-Cre mice (n=1365 cells; 2 mice). As noted above (in H), this pattern is seen to differ between *Prokr2*-Cre and *Kcnc1*-Cre lines (Χ^2^(2)=151.69, p<0.001). *Prokr2*-Cre mice showed a lower rate of cells co-expressing *Cre* and *Rbfox1* (p<0.001) as well as a lower rate of *Cre* only cells (p<0.001) that do not co-express another mRNA of interest. They also have an increased rate of *Cre* and *Vgf* co-expression (p<0.001) as well as *Cre*, *Rbfox1*, *Vgf* (p=0.005) and *Cre*, *Peg10*, and *Vgf* (p=0.010) triple expression. (**R**) Percent of cells co-expressing *Cre* and each mRNA transcript for *Prokr2*-Cre mice as a function of all cells positive for that mRNA transcript. (**S**) Relationship between *Cre* and *Peg10* mRNA estimates (n=3,909 cells) in *Prokr2*-Cre mice. (**T**) Relationship between *Cre* and *Rbfox1* mRNA estimates (n=6,217 cells) in *Prokr2*-Cre mice. (**U**) Relationship between *Cre* and *Vgf* mRNA estimates (n=8,844 cells) in *Prokr2*-Cre mice. (**V-Y**) Representative images at 40x displaying mRNA overlap of *Peg10* (*top*), *Rbfox1* (*middle*), and *Vgf* (*bottom*) with *Cre* in *Prokr2*-Cre mice. Differences in co-expression between *Kcnc1*-Cre and *Prokr2*-Cre were calculated using a Chi-squared analysis and reported Pearson residuals p-values are Bonferroni corrected.

**Supplemental Figure 3.**
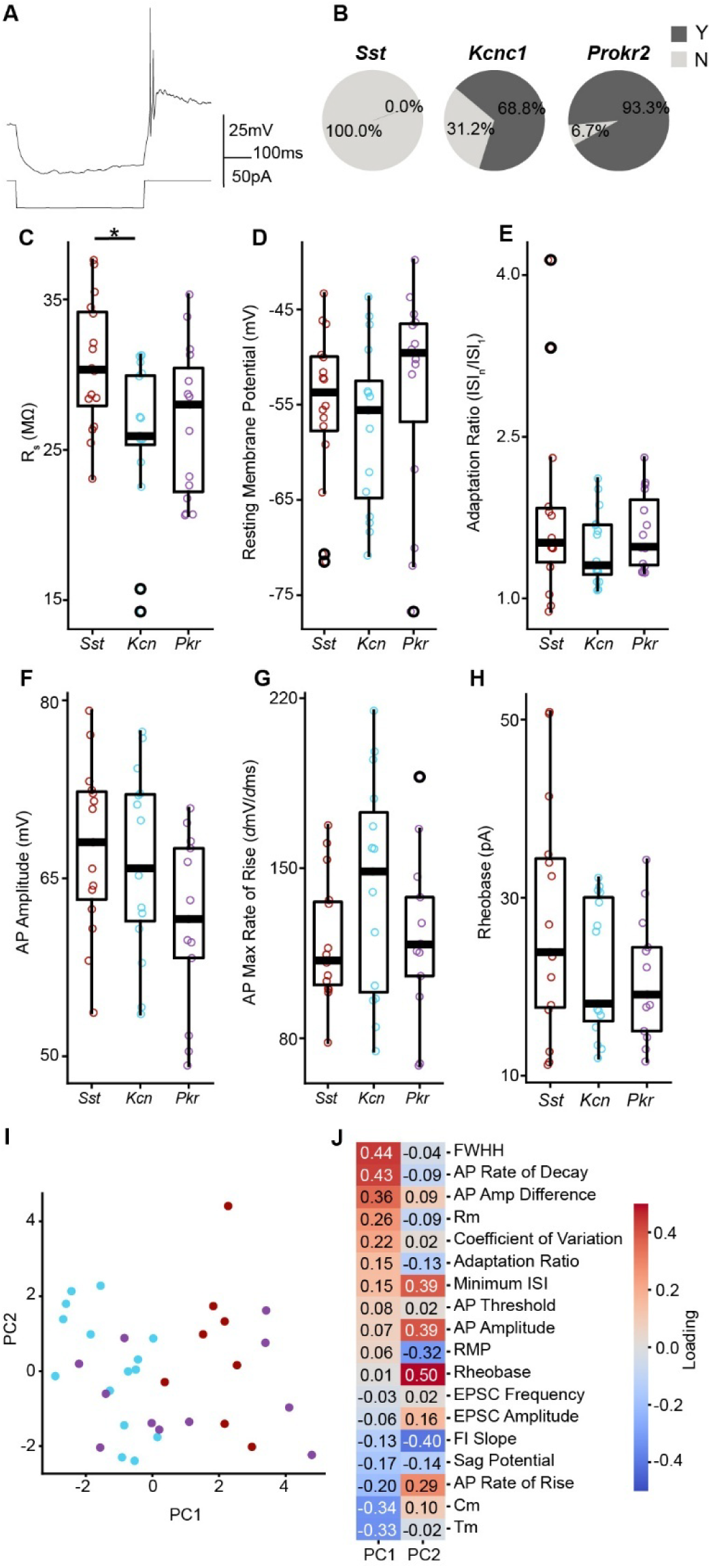
Additional LHb intrinsic electrophysiological characteristics and PCA. (**A**) Representative trace of a prolonged inhibitory rebound AP burst following a 500ms current injection of -50pA. (**B**) Percentage of cells within each transgenic mouse line that produce a single or repeated AP bursts following the injection of inhibitory current (Χ^2^(2)=29.45, p<0.001). (**C**) Average series resistance (MΩ) across recordings for each cell (F(2,42)=4.504, p=0.017) differed significantly between *Sst*+ cells and *Kcnc1*+ cells (p=0.018). (**D**) Resting membrane potential (RMP; mV) does not differ across transgenic lines (F(2,44)=0.604, p=0.551). (**E**) Adaptation ratio of interspike intervals (ms) during 500ms of 50pA current injection does not differ across transgenic lines (F(2,42)=1.289, p=0.286). (**F**) Baseline corrected amplitude (mV) of current-evoked APs does not differ across transgenic lines (F(2,40)=2.782, p=0.074). (**G**) Maximum rate of rise (*d*mV/*d*ms) from current-evoked APs does not differ across transgenic lines (F(2,40)=1.86, p=0.169). (**H**) Minimum current (pA) required to evoke an AP (rheobase) does not differ across transgenic lines (F(2,40)=1.81, p=0.177). (**I**) Principal component analysis of electrophysiological variables from *Sst*-Cre (*red*), *Kcnc1*-Cre (*blue*), and *Prokr2*-Cre (*purple*). Principal components one (PC1) and two (PC2) explain 0.412 of total variance, 0.239 and 0.173 respectively. (**J**) Variable loadings onto PC1 and PC2. Statistical analyses represent one-way ANOVAs with post-hoc Tukey’s HSD.

**Supplemental Figure 4.**
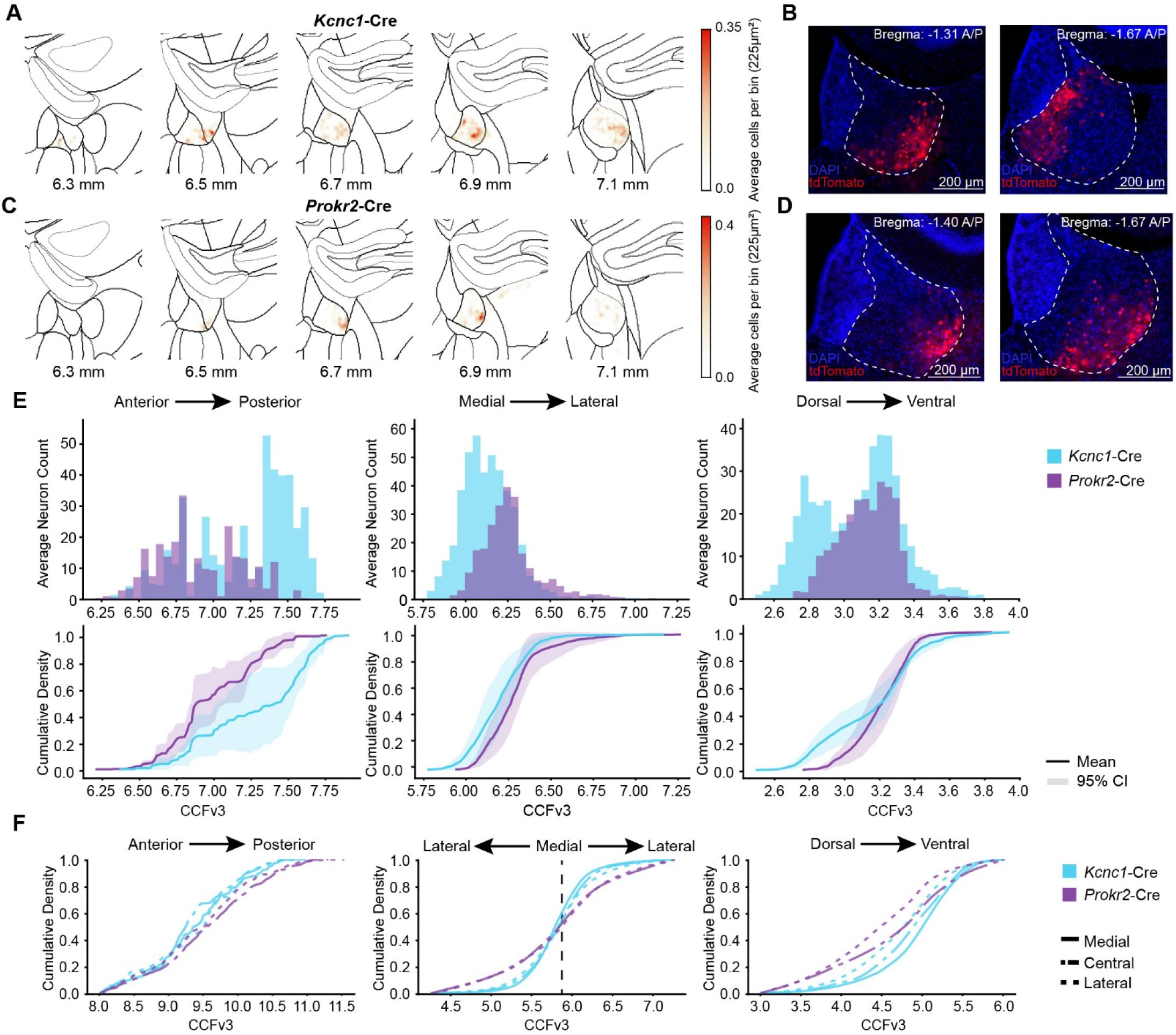
Spatial location of LHb soma for subpopulations targeted in axonal tracing. (**A**) Heatmaps representing the location of all tdTom expressing somata in *Kcnc1*-Cre animals (n=5) in 200µm sections from CCFv3 rostro-caudal position of 6300µm (approximate bregma AP: -1.0mm) to 7500µm (approximate bregma AP: -2.2mm). (**B**) Representative 20x images of intracranial injection sites for Cre-dependent tdTom expressing AAV in two *Kcnc1*-Cre animals. (**C**) Heatmaps representing the location of all tdTom expressing somata in *Prokr2*-Cre animals (n=5). (**D**) Representative 20x images of intracranial injection sites for Cre-dependent tdTom expressing AAV in two *Prokr2*-Cre animals. (**E**) Spatial density of tdTom expressing somata differs between *Kcnc1*-Cre (*blue*) and *Prokr2*-Cre (*purple*) mice across the anterior to posterior (*left;* F(9.77)=6.11, p<0.001), medial to lateral (*middle;* F(24.31)=26.88, p<0.001), and dorsal to ventral (*right;* F(18.19)=13.14, p<0.001) axes as average neuron counts across animals (top) and cumulative density (bottom). (**F**) Downstream axonal projections across the anterior to posterior (*left*), medial to lateral (*middle*), and dorsal to ventral (*right*) axes in *Kcnc1*-Cre (*blue*) and *Prokr2*-Cre (*purple*) mice for medial (*solid*), central (*dot-dashed*), and lateral (*dashed*) tdTom labeled neuronal density within the LHb. Reported statistics represent the difference in variance explained by the addition of group by position interaction to a generalized additive model.

**Supplemental Figure 5.**
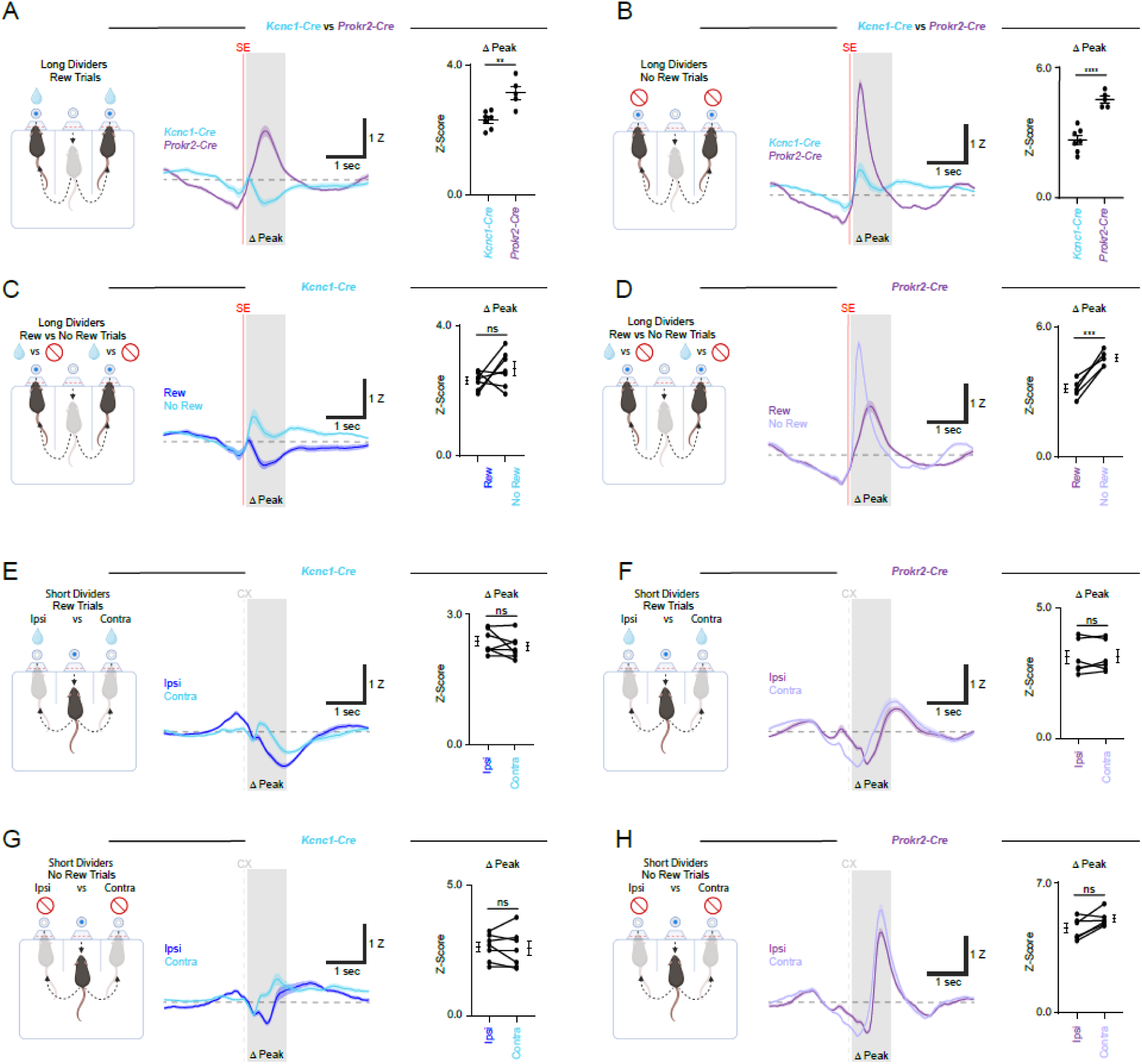
LHb subpopulation activity examining movement direction and outcome using different divider lengths. (**A – H**) Neuronal activity of *Kcnc1*-Cre and *Prokr2*-Cre mice expressing GCaMP8f. Calcium photometry data are aligned to indicated behavioral event (SE, CX, etc.). For *Kcnc1*-Cre 47 short divider and 41 long divider 2ABT sessions were combined and consists of *Kcnc1*-Cre n=7 (short and long) (4 female, 3 male) animals and for *Prokr2*-Cre 30 short divider and 25 long divider 2ABT sessions were combined across n=6 (short) or 5 (long) (2 female, 4 male for short dividers, 2 female, 3 male for long dividers) animals. For short and long dividers, *Kcnc1*-Cre consisted of 16879 and 10237 trials respectively and *Prokr2-*Cre consisted of 12145 and 9707 trials respectively. Data are Z-scored and line graphs are presented as mean (solid line) ± SEM (shaded area). Graphs quantifying Δ Peak are presented as mean ± SEM. Statistical significance is determined via two-tailed nested t-test. Diagrams to the left of data show trial type, divider type, mouse movement, and center point of line graphs. (**A**) Neuronal activity comparing *Kcnc1*-Cre vs *Prokr2*-Cre mice aligned to Side Entry on rewarded trials using long dividers. p=0.0019 (t=4.163, df=10, F=17.33). (B) Neuronal activity comparing *Kcnc1*-Cre vs *Prokr2*-Cre mice aligned to Side Entry on unrewarded trials using long dividers. ****p<0.0001 (t=6.680, df=10, F=44.62). (**C, D**) Neuronal activity aligned to Side Entry comparing rewarded to unrewarded trials using long dividers. *Kcnc1-*Cre mice. ns p=0.1409 (t=1.576, df=12, F=2.486). *Prokr2-*Cre. ***p=0.0006 (t=5.515, df=8, F=30.41). (**E, F**) Neuronal activity aligned to Center Exit comparing ipsilateral vs contralateral choice for rewarded trials using short dividers. *Kcnc1-*Cre mice. ns p=0.4513 (t=0.7786, df=12, F=0.6062). *Prokr2*-Cre. Ns p=0.9398 (t=0.07747, df=10, F=0.0066001). (**G, H**) Neuronal activity aligned to Center Exit comparing ipsilateral vs contralateral choice for rewarded trials using short dividers. *Kcnc1*-Cre mice. ns p=0.8579 (t=0.1829, df=12, F=0.03347). *Prokr2*-Cre mice ns p=0.1034 (t=1.792, df=10, F=3.210). ns=no significance. Diagrams made with Biorender.

**Supplemental Figure 6.**
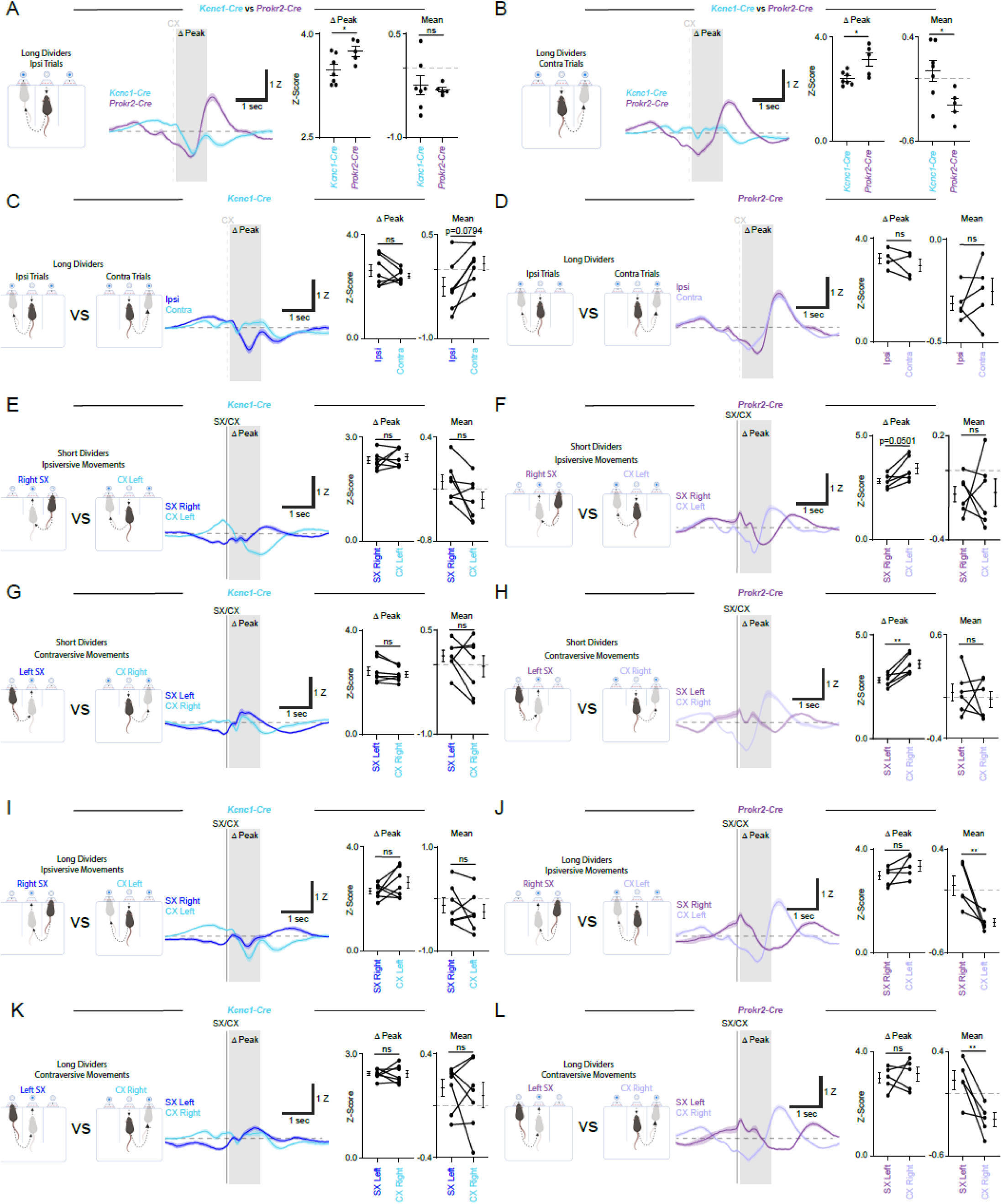
LHb activity during directional movement for LHb neuronal subpopulations. (**A – H**) Neuronal activity of *Kcnc1*-Cre and *Prokr2-*Cre mice expressing GCaMP8f. Calcium photometry data are aligned to indicated behavioral event (SE, CX, etc.). For *Kcnc1*-Cre 47 short divider and 41 long divider 2ABT sessions were combined and consists of *Kcnc1*-Cre n=7 (short and long) (4 female, 3 male) animals and for *Prokr2*-Cre 30 short divider and 25 long divider 2ABT sessions were combined across n=6 (short) or 5 (long) (2 female, 4 male for short dividers, 2 female, 3 male for long dividers) animals. For short and long dividers, *Kcnc1*-Cre consisted of 16879 and 10237 trials respectively and *Prokr2-*Cre consisted of 12145 and 9707 trials respectively. Data are Z-scored and line graphs are presented as mean (solid line) ± SEM (shaded area). Graphs quantifying Δ Peak and Mean are presented as mean ± SEM. Statistical significance is determined via two-tailed nested t-test. Diagrams to the left of data show trial type, mouse movement, and center point of line graphs. (**A**) Neuronal activity comparing *Kcnc1*-Cre vs *Prokr2*-Cre mice aligned to Center Exit for trials to ipsilateral port using long dividers. Δ Peak *p=0.0379 (t=2.392, df=10, F=5.720), Mean ns p=0.7144 (t=0.3765, df=10, F=0.1418). (**B**) Neuronal activity comparing *Kcnc1*-Cre vs *Prokr2-*Cre mice aligned to Center Exit for trials to the contralateral port using long dividers. Δ Peak *p=0.0149 (t=2.934, df=10, F=8.609), Mean *p=0.0309 (t=2.511, df=10, F=6.304). (**C, D**) Neuronal activity aligned to Center Exit comparing trials to the ipsilateral port with those to the contralateral port using long dividers. *Kcnc1*-Cre mice Δ Peak ns p=0.4250 (t=0.8258, df=12, F=0.6819), Mean ns p=0.0794 (t=1.917, df=12, F=3.675). *Prokr2*-Cre mice Δ Peak ns p=0.5618 (t=0.6053, df=8, F=0.3664), Mean ns p=0.4551 (t=0.7849, df=8, F=0.6160). (**E, F**) Neuronal activity comparing ipsiversive movements (those trials moving ipsilaterally regardless of port) aligned to right port Side Exit vs. Center Exit to the left using short dividers. *Kcnc1*-Cre mice Δ Peak ns p=0.4856 (t=0.7195, df=12, F=5.177), Mean ns p=0.1172 (t=1.688, df=12, F=2.849). *Prokr2*-Cre mice Δ Peak ns p=0.0501 (t=2.227, df=10, F=4.961), Mean ns p=0.9190 (t=0.1043, df=10, F=0.01088). (**G, H**) Neuronal activity comparing contraversive movements (those trials moving contralaterally regardless of port) aligned to left port Side Exit vs. Center Exit to the right using short dividers. *Kcnc1*-Cre mice Δ Peak ns p=0.5420 (t=0.6276, df=12, F=0.3938), Mean ns p=0.4008 (t=0.8710, df=12, F=0.7587). *Prokr2-*Cre mice Δ Peak **p=0.0083 (t=3.281, df=10, F=10.76), Mean ns p=0.5612 (t=0.6010, df=10, F=0.3612). (**I, J**) Neuronal activity comparing ipsiversive movements (those trials moving ipsilaterally regardless of port) aligned to right port Side Exit vs. Center Exit to the left using long dividers. *Kcnc1*-Cre mice Δ Peak ns p=0.2121 (t=1.318, df=12, F=1.738), Mean ns p=0.5238 (t=0.6566, df=12, F=0.4311). *Prokr2*-Cre mice Δ Peak ns p=0.1799 (t=1.469, df=8, F=2.159), Mean **p=0.0077 (t=3.530, df=8, F=12.46). (**K, L**) Neuronal activity comparing contraversive movements (those trials moving contralaterally regardless of port) aligned to left port Side Exit vs. Center Exit to the right using long dividers. *Kcnc1*-Cre mice Δ Peak ns p=0.9876 (t=0.01587, df=12, F=0.0002517), Mean ns p=0.6471 (t=0.4695, df=12, F=0.2204). *Prokr2*-Cre mice Δ Peak ns p=0.6636 (t=0.4516, df=8, F=0.2039), Mean **p=0.0096 (t=3.381, df=8, F=11.43). ns=no significance. Diagrams made with Biorender.

**Supplemental Figure 7.**
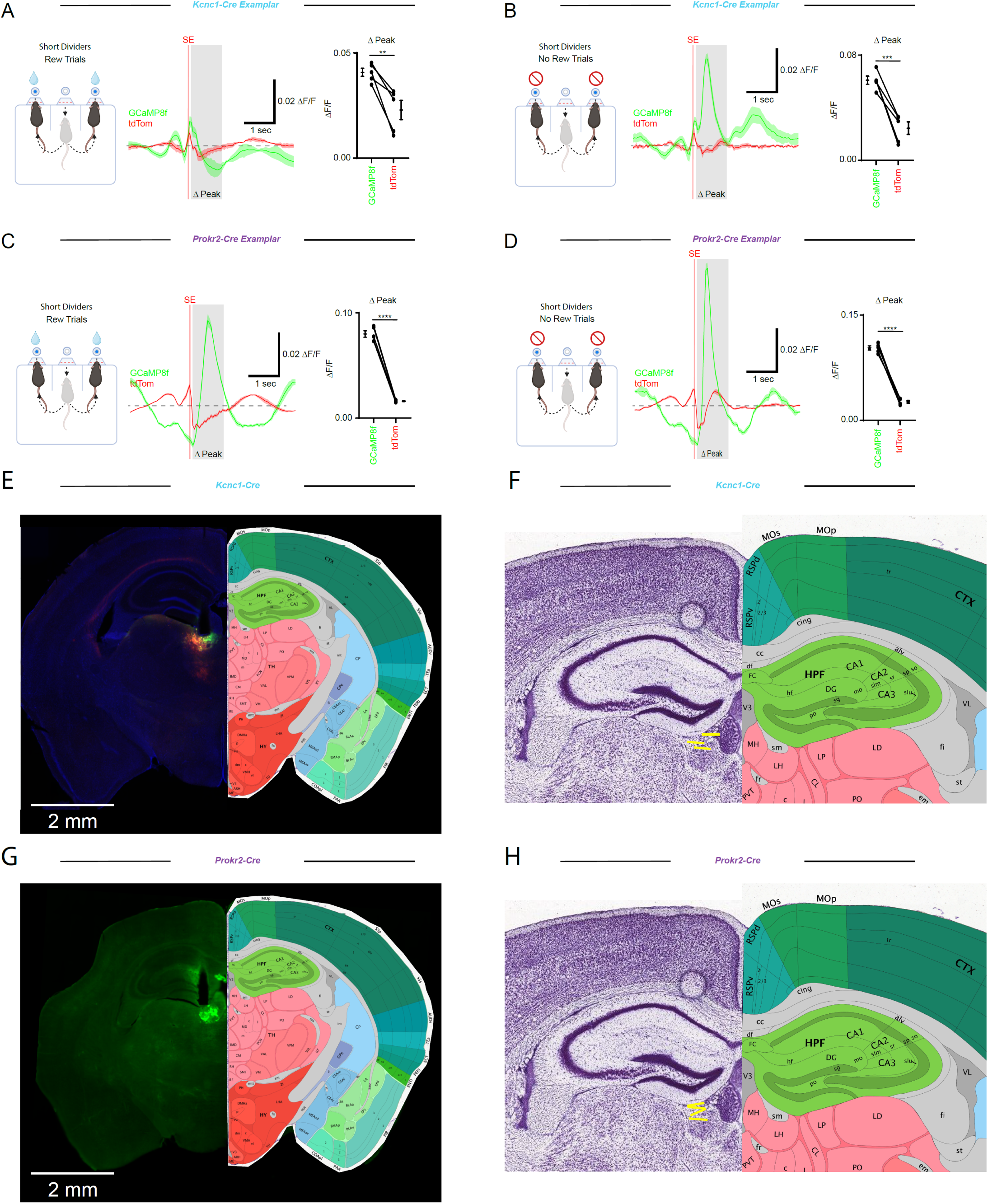
Control fluorophore fluorescence recorded simultaneously with GCaMP8f fluorescence and viral expression/ fiber placement anatomy. Neuronal activity of *Kcnc1*-Cre mice expressing Cre-dependent GCaMP8f and Cre-dependent tdTom. Data includes multiple daily 2ABT sessions that are combined consisting of *Kcnc1*-Cre n=3 (2 female, 1 male). Data are Z-scored and line graphs are presented as mean (solid line) ± SEM (shaded area). Graphs quantifying Δ Peak are presented as mean ± SEM. Statistical significance is determined via two-tailed nested t-test. Diagrams to the left of data show trial type, mouse movement, and center point of line graphs (*Kcnc1*-Cre n=3 (2 female, 1 male) and 7898 trials). (**A**) Neuronal activity comparing green and red fluorescence in *Kcnc1*-Cre mice aligned to Side Entry on rewarded trials using short dividers. *p<0.05 (t=4.079, df=4, F=16.64). (**B**) Neuronal activity comparing green and red fluorescence in *Kcnc1*-Cre mice aligned to Side Entry on unrewarded trials using short dividers. p=0.0548 (t=2.687, df=4, F=7.222). (**C**) Neuronal activity comparing green and red fluorescence in *Kcnc1*-Cre mice aligned to Center Exit for all ipsilateral trials using short dividers. *p<0.05 (t=4.026, df=4, F=16.21). (**D**) Neuronal activity comparing green and red fluorescence in *Kcnc1*-Cre mice aligned to Center Exit for all contralateral trials using short dividers. ns=no significance (t=2.224, df=4, F=4.948). (**E**) Example brain slice showing GCaMP8f and tdTom expression in *Kcnc1*+ neurons in the LHb as well as probe tip location (DAPI shown in blue). (**F**) Allen Brain Institute brain image and atlas with probe locations (yellow line) of all photometry mice (n=3). (**G**) Example brain slice showing GCaMP8f expression in *Prokr2*+ neurons in the LHb as well as probe tip location. (H) Allen Brain Institute brain image and atlas with probe locations (yellow line) of all photometry mice (n=4). Diagrams made with Biorender.

**Supplemental Figure 8.**
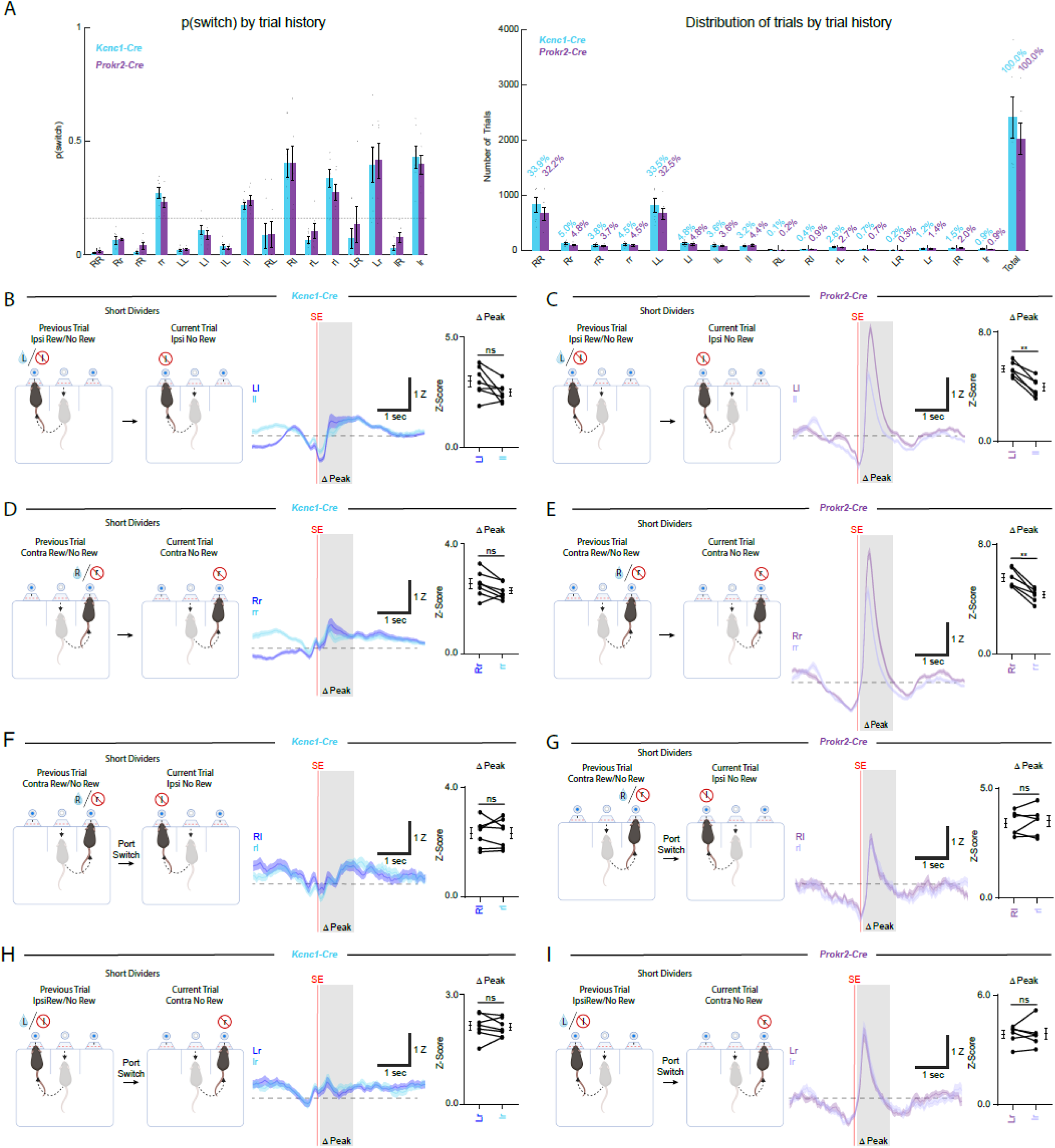
The effects of laterality on trial history for *Prokr2*+ and *Kcnc1*+ neuronal activity during the 2ABT. (**A**) The distribution of different trial histories across all trials (percentages of total trials denoted above each trial type) (*left)*. Trial history dependence of p(switch) for *Kcnc1*-Cre and *Prokr2*-Cre mice, dashed gray line is mean p(switch) for all trials (*right*) (bar represents mean, error bar represents ±SEM. (**B – I**) Neuronal activity of *Kcnc1*-Cre and *Prokr2*-Cre mice expressing GCaMP8f. For *Kcnc1*-Cre 47 short divider 2ABT sessions were combined and consists of *Kcnc1*-Cre n=7 (4 female, 3 male) animals and for *Prokr2*-Cre 30 short divider 2ABT sessions were combined across n=6 (2 female, 4 male) animals. *Kcnc1*-Cre mice consisted of 16879 trials and *Prokr2-*Cre mice consisted of 12145 trials. Data are Z-scored and line graphs are presented as mean (solid line) ± SEM (shaded area). Graphs quantifying Δ Peak are presented as mean ± SEM. Statistical significance is determined via two-tailed nested t-test. Diagrams to the left of data show trial type, mouse movement, and center point of line graphs. (**B, C**) Neuronal activity aligned to Side Entry comparing unrewarded trials to the left (ipsilateral) port where the previous trial was either rewarded or unrewarded to the same ipsilateral port using short dividers. *Kcnc1*-Cre mice. ns p=0.1200 (t=1.674, df=12, F=2.802). *Prokr2*-Cre mice. **p=0.0038 (t=3.744, df=10, F=14.02). (**D, E**) Neuronal activity aligned to Side Entry comparing unrewarded trials to the right (contralateral) port where the previous trial was either rewarded or unrewarded to the same contralateral port using short dividers. *Kcnc1*-Cre mice. ns p=0.2197 (t=1.295, df=12, F=1.677). *Prokr2*-Cre mice. **p=0.0041 (t=3.708, df=10, F=13.75). (**F, G**) Neuronal activity aligned to Side Entry comparing unrewarded trials to the left (ipsilateral port) where the previous trial was either rewarded or unrewarded to the opposite contralateral port using short dividers. *Kcnc1*-Cre mice. ns p=0.9641 (t=0.04597, df=12, F=0.002114). *Prokr2*-Cre mice. ns p=0.7896 (t=0.2740, df=10, F=0.07509). (**H, I**) Neuronal activity aligned to Side Entry comparing unrewarded trials to the right (contralateral) port where the previous trial was either rewarded or unrewarded to the opposite ipsilateral port using short dividers. *Kcnc1*-Cre mice. ns p=0.8004 (t=0.2585, df=12, F=0.06684). *Prokr2*-Cre mice. ns p=0.8899 (t=0.1420, df=10, F=0.02017). ns=no significance. Diagrams made with Biorender.

